# The Sociodemographic and Lifestyle Correlates of Epigenetic Aging in a Nationally Representative U.S. Study of Younger Adults

**DOI:** 10.1101/2024.03.21.585983

**Authors:** Kathleen Mullan Harris, Brandt Levitt, Lauren Gaydosh, Chantel Martin, Jess M. Meyer, Aura Ankita Mishra, Audrey L. Kelly, Allison E. Aiello

**Affiliations:** Department of Sociology, University of North Carolina at Chapel Hill, Chapel Hill, NC; Carolina Population Center, University of North Carolina at Chapel Hill, Chapel Hill, NC; Department of Sociology, University of Texas at Austin, Austin, TX; Population Research Center, University of Texas at Austin, Austin, TX; Department of Epidemiology, Gillings School of Global Public Health, University of North Carolina at Chapel Hill, Chapel Hill, NC; Department of Population Health, University of Kansas Medical Center, Kansas City, KS; Department of Psychology, North Carolina State University, Raleigh, NC; Department of Epidemiology, Mailman School of Public Health, Columbia University, New York, NY; Robert N. Butler Columbia Aging Center, Mailman School of Public Health, Columbia University, New York, NY

## Abstract

**Importance:** Epigenetic clocks represent molecular evidence of disease risk and aging processes and have been used to identify how social and lifestyle characteristics are associated with accelerated biological aging. However, most of this research is based on older adult samples who already have measurable chronic disease.

**Objective:** To investigate whether and how sociodemographic and lifestyle characteristics are related to biological aging in a younger adult sample across a wide array of epigenetic clock measures.

**Design:** Nationally representative prospective cohort study.

**Setting:** United States (U.S.).

**Participants:** Data come from the National Longitudinal Study of Adolescent to Adult Health, a national cohort of adolescents in grades 7-12 in U.S. in 1994 followed for 25 years over five interview waves. Our analytic sample includes participants followed-up through Wave V in 2016-18 who provided blood samples for DNA methylation (DNAm) testing (n=4237) at Wave V.

**Exposure:** Sociodemographic (sex, race/ethnicity, immigrant status, socioeconomic status, geographic location) and lifestyle (obesity status, exercise, tobacco, and alcohol use) characteristics.

**Main Outcome:** Biological aging assessed from blood DNAm using 16 epigenetic clocks when the cohort was aged 33-44 in Wave V.

**Results:** While there is considerable variation in the mean and distribution of epigenetic clock estimates and in the correlations among the clocks, we found sociodemographic and lifestyle factors are more often associated with biological aging in clocks trained to predict current or dynamic phenotypes (e.g., PhenoAge, GrimAge and DunedinPACE) as opposed to clocks trained to predict chronological age alone (e.g., Horvath). Consistent and strong associations of faster biological aging were found for those with lower levels of education and income, and those with severe obesity, no weekly exercise, and tobacco use.

**Conclusions and Relevance:** Our study found important social and lifestyle factors associated with biological aging in a nationally representative cohort of younger-aged adults. These findings indicate that molecular processes underlying disease risk can be identified in adults entering midlife before disease is manifest and represent useful targets for interventions to reduce social inequalities in heathy aging and longevity.

**Key Points:** *Question:* Are epigenetic clocks, measures of biological aging developed mainly on older-adult samples, meaningful for younger adults and associated with sociodemographic and lifestyle characteristics in expected patterns found in prior aging research?

*Findings:* Sociodemographic and lifestyle factors were associated with biological aging in clocks trained to predict morbidity and mortality showing accelerated aging among those with lower levels of education and income, and those with severe obesity, no weekly exercise, and tobacco use.

*Meaning:* Age-related molecular processes can be identified in younger-aged adults before disease manifests and represent potential interventions to reduce social inequalities in heathy aging and longevity.

## Introduction

Research on aging has long documented social and demographic differentials in morbidity and mortality risk, but only recently has been able to explore the underlying molecular and cellular changes that accompany aging processes and shorten life. New developments in geroscience have identified biological “hallmarks of aging”^1–3^ about which studies now collect data to examine the social determinants of biological aging.^4–8^ Epigenomic profiling, in particular, has greatly increased the availability of novel indicators of biological aging in the form of epigenetic clocks, ^9–11^ composite measures of DNA methylation (DNAm) that represent molecular evidence of disease risk and aging processes.^12,13^ Epigenetic clocks predict epigenetic age, and the relative comparison of epigenetic age with chronological age represents epigenetic age acceleration (EAA), providing a measurement of differences in biological aging among individuals of the same calendar age.^14^ Hereafter we refer to EAA as biological aging. Epigenetic clock measures have been linked to many age-related diseases and mortality.^15–26^ Thus, they potentially represent useful targets for interventions to reduce social inequalities in healthy aging and longevity^26^, particularly if the clocks can detect biological aging in young individuals without apparent disease.

Considerable progress has been made in measuring biological age using numerous sources of data to create epigenetic clocks.^7,27,28^ First-generation clocks were constructed as molecular predictors of chronological age.^29–31^ Attention soon shifted, however, to predict aging outcomes above and beyond chronological age. Thus, second-generation epigenetic clocks were calibrated on differences in biological aging reflected by disease and mortality risks.^20,32^ Recent advances exploit longitudinal measurement of aging phenotypes to construct clocks that capture the pace of change in biological aging over time,^33,34^ representing a third generation of epigenetic clocks.

There is a growing body of research on the social and lifestyle factors associated with biological aging as measured using EAA^7,28^ providing insights for differential exposures that influence aging.^35–39^ To the extent that certain social and lifestyle factors are known to be associated with greater age-related health risks, we would expect these factors to be associated with more rapid biological aging, affirming the research value of epigenetic clocks as markers of aging, particularly in younger adults. Indeed, lower socioeconomic status in education, income, and wealth has been associated with more rapid epigenetic aging.^8,40^ Variability in biological aging by sex generally shows males experience more rapid epigenetic aging,^28,39,41^ whereas variability by race/ethnicity is inconsistent, with positive, negative, and null differences observed in previous research comparing Black or Hispanic to White individuals.^5,9^ Among lifestyle factors, existing research shows strong and consistently positive associations between tobacco use and BMI with biological aging in second- and third-generation clocks, but more mixed results for alcohol use and exercise.^7–9,28^ Overall, results suggest that more recent second- and third-generation clocks are more sensitive to social and environmental exposures, though more work is needed to better understand whether and how clocks capture shared or distinct aspects of aging.

Existing research examining sociodemographic factors and biological aging has notable gaps. First, many studies use only a single or a few clocks, making it difficult to ascertain whether the results are consistent across clocks. Second, most epigenetic research relies on small, often local, and non-diverse samples, limiting generalizability.^7,1413,40^ Some recent studies have used large representative samples that examined a range of clocks, but focused on older adults.^10,41^ In fact, the majority of research on epigenetic aging has been based on samples of older adults^10,1415,20,22–24,38,42–45^ (exceptions include^46,47^). Given that older adults are more likely to have chronic comorbidities, it is difficult to disentangle the impacts of underlying disease from sociodemographic exposures. Further, as age increases, biological age may become a less reliable predictor of health outcomes due to mortality selection and increased biological heterogeneity in older age.

Our research addresses the existing gaps by investigating the relationship between sociodemographic and lifestyle factors and 16 DNAm measures of biological aging in a diverse population of adults aged 33-44 from the U.S. representative National Longitudinal Study of Adolescent to Adult Health (Add Health). Using new methylation data with national representation of racial, ethnic, socioeconomic, and geographic groups, we contribute to the limited research on epigenetic aging in younger adults. Our study is one of the few to examine the emergence of sociodemographic inequalities in aging before adults enter midlife across established epigenetic clock measures.

## Methods

Data come from Add Health, a nationally representative cohort study of U.S. adolescents in grades 7-12 in 1994 and followed for 25 years across five interview waves.^42^ We use data primarily from Waves I (1994-95) and V (2016-18) when the cohort was aged 33-44. During the Wave V survey, 5,381 participants consented and were scheduled for a follow-up in-person home exam when venous blood was drawn (94% consent rate) for DNAm assay. After removing samples that did not pass quality control and eliminating replicates, the DNAm sample included 4,582 participants. The sample size is further reduced to 4,237 due to missing values on social and demographic factors.

Methylation analysis was conducted using the Illumina Infinium chemistry.^43^ DNAm levels across ∼850,000 CpG sites were measured using the Infinium Methylation EPIC BeadChip (Illumina, Inc.; San Diego, CA). Beta values for CpG sites across the genome were measured according to kit protocols and filtered to remove polymorphic positions. Remaining CpG sites were restricted to a set of 30,484 CpG sites used with the DNA methylation calculator (https://dnamage.genetics.ucla.edu) and Methylcipher^44^ and Dunedin^33,34^ calculators; PC clocks based on 78,464 CpG sites and used code publicly available on GitHub (see **Supplement 1 for data quality and curation**).

### Epigenetic Clocks

We constructed 16 epigenetic clocks when the cohort was average age 38.4 at the time of venous blood draw (**Table 1; details in Supplement 2**). Clocks are shown in order of their “generation” beginning with five first-generation clocks: Horvath1, Horvath2, Hannum, VidalBralo, and Zhang2019. The two Horvath clocks were constructed across multiple tissues as a pan-tissue clock of chronological age and differ according to the number of CpG sites on which the algorithm was based.^30,31^ The Hannum^29^ and VidalBralo^45^ clocks were trained on blood samples and Zhang2019^46^ was trained on blood and saliva samples to predict chronological age. We examined three second-generation clocks that were trained on disease phenotypes and mortality in prediction of epigenetic age including Lin,^47^ PhenoAge,^20^ and GrimAge^21^—which also incorporated smoking-related methylation changes.

**Table 1.** Weighed Descriptive Statistics for DNA Methylation Epigenetic Clocks and Pearson Correlation with Age^a^ (N= 4237).

| Type of Measure | Measure | Mean | SD | Min | Max | Pearson R | Pearson Prob |
| --- | --- | --- | --- | --- | --- | --- | --- |
| Chronological Age<sup>a</sup> | Years of age | 38.4 | 2.0 | 33.1 | 44.8 | 1.00 | 0.00 |
| Clocks | Horvath1 | 39.1 | 4.3 | 15.5 | 62.3 | 0.42 | 0.00 |
|  | Horvath2 | 35.4 | 3.5 | 17.7 | 57.4 | 0.58 | 0.00 |
|  | Hannum | 31.6 | 3.9 | 13.2 | 47.5 | 0.40 | 0.00 |
|  | VidalBralo | 54.9 | 3.5 | 38.2 | 77.6 | 0.27 | 0.00 |
|  | Zhang2019 | 32.0 | 4.1 | 19.9 | 58.3 | 0.62 | 0.00 |
|  | Lin | 23.1 | 5.2 | 4.0 | 44.0 | 0.32 | 0.00 |
|  | PhenoAge | 30.1 | 5.7 | 12.8 | 50.9 | 0.32 | 0.00 |
|  | GrimAge | 52.5 | 4.6 | 39.1 | 71.9 | 0.39 | 0.00 |
| PC Clocks | PCHorvath1 | 45.3 | 3.7 | 31.6 | 66.0 | 0.36 | 0.00 |
|  | PCHorvath2 | 41.2 | 4.1 | 26.6 | 62.5 | 0.32 | 0.00 |
|  | PCHannum | 46.1 | 3.8 | 31.5 | 64.8 | 0.38 | 0.00 |
|  | PCPhenoAge | 41.0 | 5.3 | 21.7 | 66.4 | 0.34 | 0.00 |
|  | PCGrimAge | 54.6 | 3.9 | 43.6 | 71.3 | 0.43 | 0.00 |
| Age Acceleration | Horvath1AA | 0.1 | 5.5 | -25.1 | 26.0 | 0.00 | 0.78 |
|  | Horvath2AA | 0.0 | 3.9 | -15.9 | 21.8 | -0.01 | 0.52 |
|  | HannumAA | 0.1 | 5.0 | -25.5 | 25.0 | 0.00 | 0.93 |
|  | VidalBraloAA | -0.1 | 4.8 | -23.0 | 24.0 | 0.00 | 0.94 |
|  | Zhang2019AA | 0.0 | 4.4 | -18.2 | 26.2 | -0.02 | 0.28 |
|  | LinAA | 0.1 | 7.2 | -30.1 | 31.8 | -0.01 | 0.65 |
|  | PhenoAgeAA | 0.1 | 7.8 | -32.4 | 29.9 | 0.01 | 0.66 |
|  | GrimAgeAA | 0.3 | 6.2 | -15.7 | 26.6 | 0.01 | 0.54 |
| PC Age Acceleration | PCHorvath1AA | 0.1 | 5.0 | -17.9 | 31.8 | 0.00 | 0.97 |
|  | PCHorvath2AA | 0.0 | 5.5 | -20.7 | 26.9 | 0.00 | 0.90 |
|  | PCHannumAA | 0.1 | 5.0 | -24.4 | 29.8 | -0.01 | 0.67 |
|  | PCPhenoAgeAA | 0.1 | 7.2 | -29.1 | 32.8 | 0.00 | 0.89 |
|  | PCGrimAgeAA | 0.3 | 5.2 | -14.5 | 25.4 | 0.00 | 0.78 |
| Rate of Aging | Dunedin PoAm | 0.0 | 1.0 | -3.9 | 4.9 | -0.01 | 0.52 |
|  | Dunedin PACE | 0.0 | 1.0 | -4.5 | 4.6 | 0.08 | 0.00 |
| Risk of mortality | Zhang2017 | -1.3 | 0.4 | -2.6 | 0.4 | 0.17 | 0.00 |

<sup>a</sup> Chronological age at Wave V blood draw.

To reduce technical variation in CpG beta values on which epigenetic clocks are based, Higgins-Chen and colleagues^48^ retrained the Hannum, Horvath1, Horvath2, GrimAge, and PhenoAge clocks on principal components (PCs) of CpG methylation values rather than on individual CpGs with the goal of reducing the effects of technical noise at any given individual CpG. We refer to these as PC clocks (**Table 1**). For each of these 13 first- and second-generation clocks, we calculate EAA by taking the residuals of the clock values regressed on chronological age.

The final set of three clocks we refer to as third generation. Their biological aging estimates use a different unit of measurement: DunedinPoAm, Dunedin PACE, and Zhang2017. DunedinPoAm^33^ and Dunedin PACE^34^ estimate the pace of biological aging in standard deviation (SD) units based on change in biomarkers of organ system dysfunction and Zhang2017 estimates a continuous risk score of all-cause mortality.^49^

### Sociodemographic and Lifestyle Characteristics

Social and demographic factors include a continuous measure of chronological age at blood draw and categorical measures of sex assigned at birth (female, male); race/ethnicity (White, Black, Hispanic, Asian/Pacific Islander, and other); immigrant generation (first, second, third+); education (college+, some college, no college); household income (>100k, 50-100k, 25-50k, <25k); region of residence (Northeast, West, Midwest, South); and rural/urban residence (metropolitan, micropolitan, small town/rural). Lifestyle measures include obesity status (normal or underweight [BMI<25], overweight [25≤BMI<30], obese [30≤BMI<40], severely obese [BMI≥40]); bouts of moderate to rigorous exercise per week (5+, 1-4, 0); tobacco use (never, former, current); and alcohol use (none, light, heavy/binge). See **Supplement 2** for additional details on variable construction.

### Analysis Plan

We conduct descriptive statistical analysis on the 16 clocks and sample characteristics and examine correlations among the epigenetic clocks, chronological age, and measures of accelerated and pace of aging. We perform weighted linear regression of accelerated biological aging on sociodemographic and lifestyle factors both in bivariate and multivariable models (with and without controls for cell composition) to assess their independent associations with biological aging using the study survey weights and the R statistical library “Survey”.^50,51^

## Results

The weighted mean and range of epigenetic ages for the various clocks (**Table 1**) show considerable variation from a mean of 23.1 years (Lin) to 54.9 (VidalBralo) for the cohort (average chronological age 38.4). Estimates of epigenetic age are positively correlated with chronological age (**Table 1**). In general, the first-generation clocks are more strongly related with age (except for VidalBralo). Second-generation clocks and PC clocks have moderate correlations with chronological age. While DunedinPoAm is not correlated with age, DunedinPACE and the Zhang2017 risk of mortality measure have small and significant correlations with age.

Sociodemographic and lifestyle characteristics of the sample (**Table 2**) reflect national representation of sex (51% female), racial and ethnic groups (71% White; 16.7% Black; 8.7% Hispanic; 3.5% Asian/Pacific Islander/other), and immigrant status (14.4% are first or second generation). The majority have at least some college education (83%) and one-third have household incomes above $100,000. Reflecting current health trends, obesity (33%) and severe obesity (10%) levels were high, but only 23% get no exercise during the week, 26% are current smokers, and 47% engage in heavy or binge drinking.

**Table 2.** Descriptive Statistics of Sample Characteristics (unweighted N and weighted %, N=4237).

| **Variable** | **N** | **Percentage** |
| --- | --- | --- |
| Sex |  |  |
| Female | 2569 | 51.30 |
| Male | 1668 | 48.70 |
| Race/Ethnicity |  |  |
| White | 2746 | 71.00 |
| Black | 811 | 16.70 |
| Hispanic | 435 | 8.70 |
| Asian/Pacific | 218 | 2.70 |
| Other | 27 | 0.80 |
| Immigrant Generation |  |  |
| First | 200 | 4.40 |
| Second | 551 | 10.00 |
| Third+ | 3486 | 85.70 |
| Education |  |  |
| College+ | 2005 | 42.70 |
| Some college | 1622 | 40.00 |
| No college | 610 | 17.30 |
| Income |  |  |
| >100k | 1491 | 32.50 |
| 50-100k | 1403 | 33.60 |
| 25-50k | 743 | 18.40 |
| <25k | 600 | 15.50 |
| Region of Residence |  |  |
| Northeast | 438 | 9.60 |
| West | 991 | 17.70 |
| Midwest | 1162 | 31.20 |
| South | 1646 | 41.40 |
| Rural/Urban residence |  |  |
| Metropolitan | 3711 | 85.90 |
| Micropolitan | 286 | 7.90 |
| Small town/rural | 240 | 6.10 |
| Obesity |  |  |
| Normal or underweight | 1173 | 26.00 |
| Overweight | 1304 | 31.50 |
| Obese | 1334 | 32.70 |
| Severe obesity | 426 | 9.90 |
| Moderate to Rigorous Exercise |  |  |
| 5 or more bouts/week | 1612 | 38.10 |
| 1-4 bouts/week | 1656 | 39.30 |
| No bouts/week | 969 | 22.60 |
| Tobacco |  |  |
| Never | 2429 | 52.30 |
| Former | 817 | 21.90 |
| Current | 991 | 25.90 |
| Alcohol |  |  |
| None | 231 | 6.08 |
| Light | 2035 | 47.11 |
| Heavy/Binge | 1971 | 46.81 |

All clocks were positively correlated with each other (**Figure 1A**), with the strongest correlations between the first-generation clocks (except for VidalBralo) and especially among the PC versions of these clocks. Among the second-generation clocks, PhenoAge had a more consistent positive correlation with the other clocks. Correlations among the age acceleration measures were somewhat lower and less likely to reach statistical significance (**Figure 1B**). In general, correlations between age acceleration measures are stronger within generations of clocks compared to across generations. The one exception is PhenoAgeAA, which was consistently correlated with all generations of clocks.

**Figure 1:**
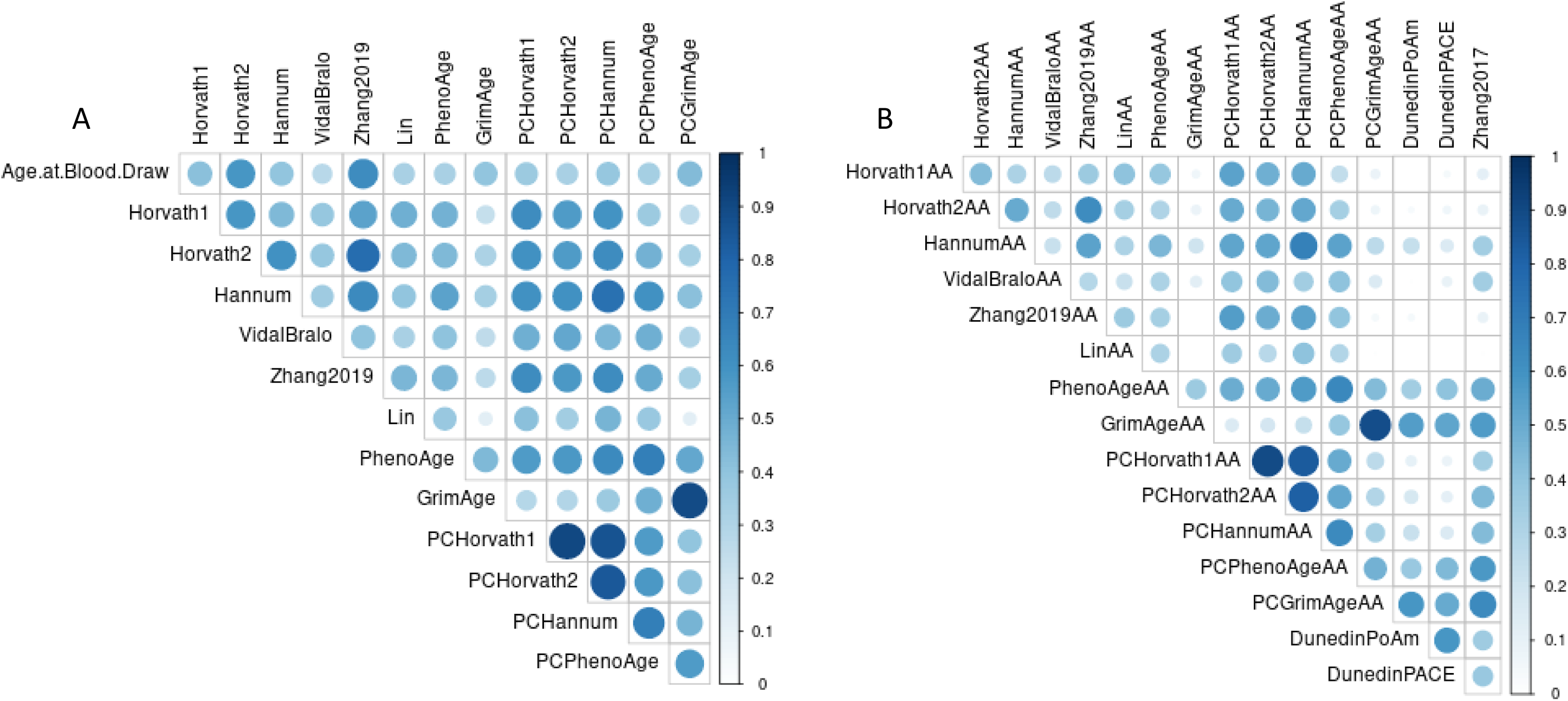
Correlations between epigenetic age clocks (A) and age acceleration and pace of aging measures (B). Notes: Darker shade and larger circle indicates greater strength of the correlation estimate. Blank squares indicate correlation not significant.

Significant bivariate associations between sociodemographic and lifestyle factors and biological aging measures (**Figure 2; see Supplement Table 1**) were more prominent among the second-generation clocks PhenoAgeAA, GrimAgeAA, PCPhenoAgeAA, and PCGrimAgeAA and third-generation DunedinPoAm and DunedinPACE than first-generation. LinAA had the fewest associations among second-generation clocks. Social and lifestyle factors were highly related to third-generation clocks (DunedinPoAm, DunedinPACE, Zhang2017) shown in standard deviation and risk units.

**Figure 2.**
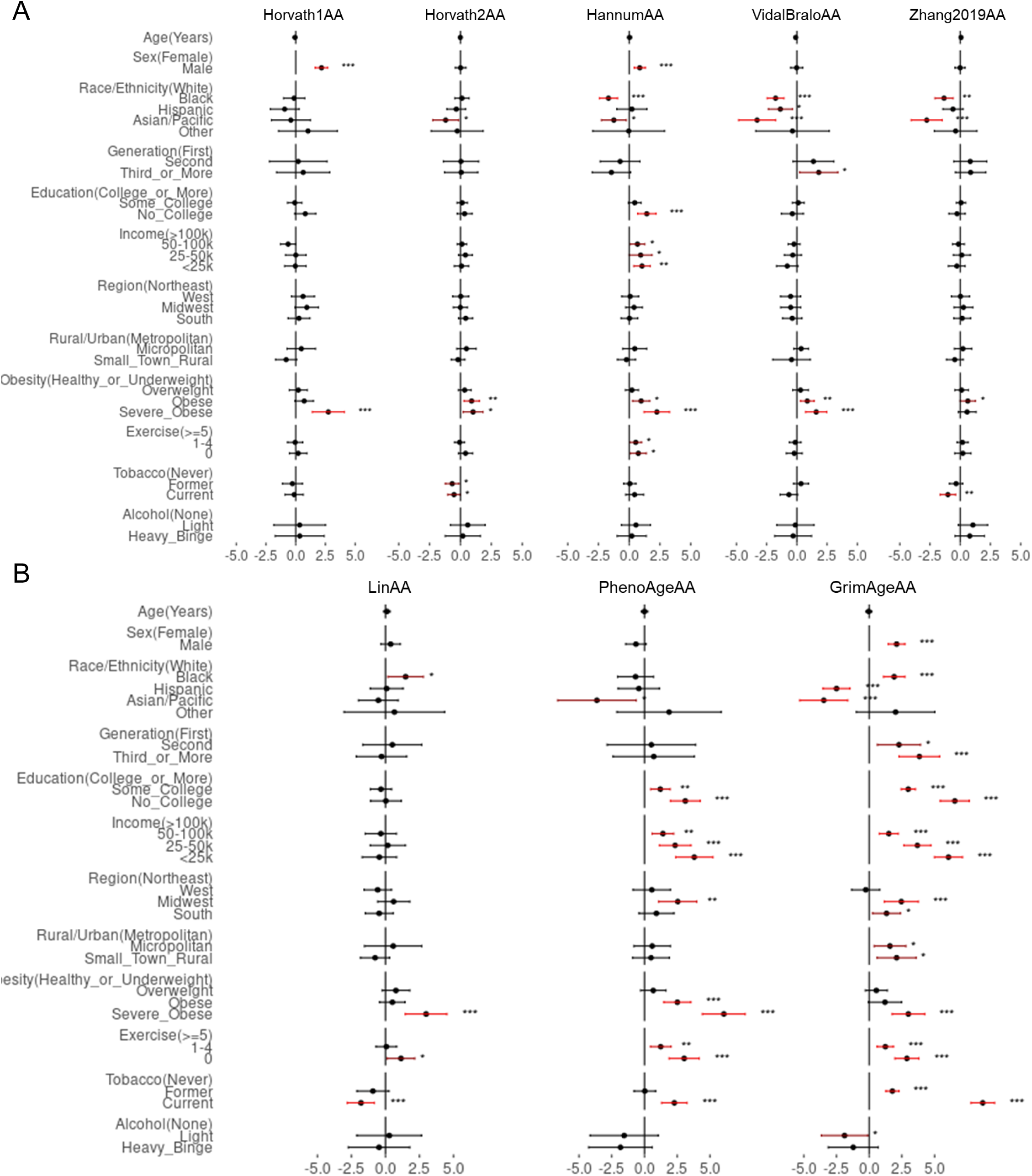
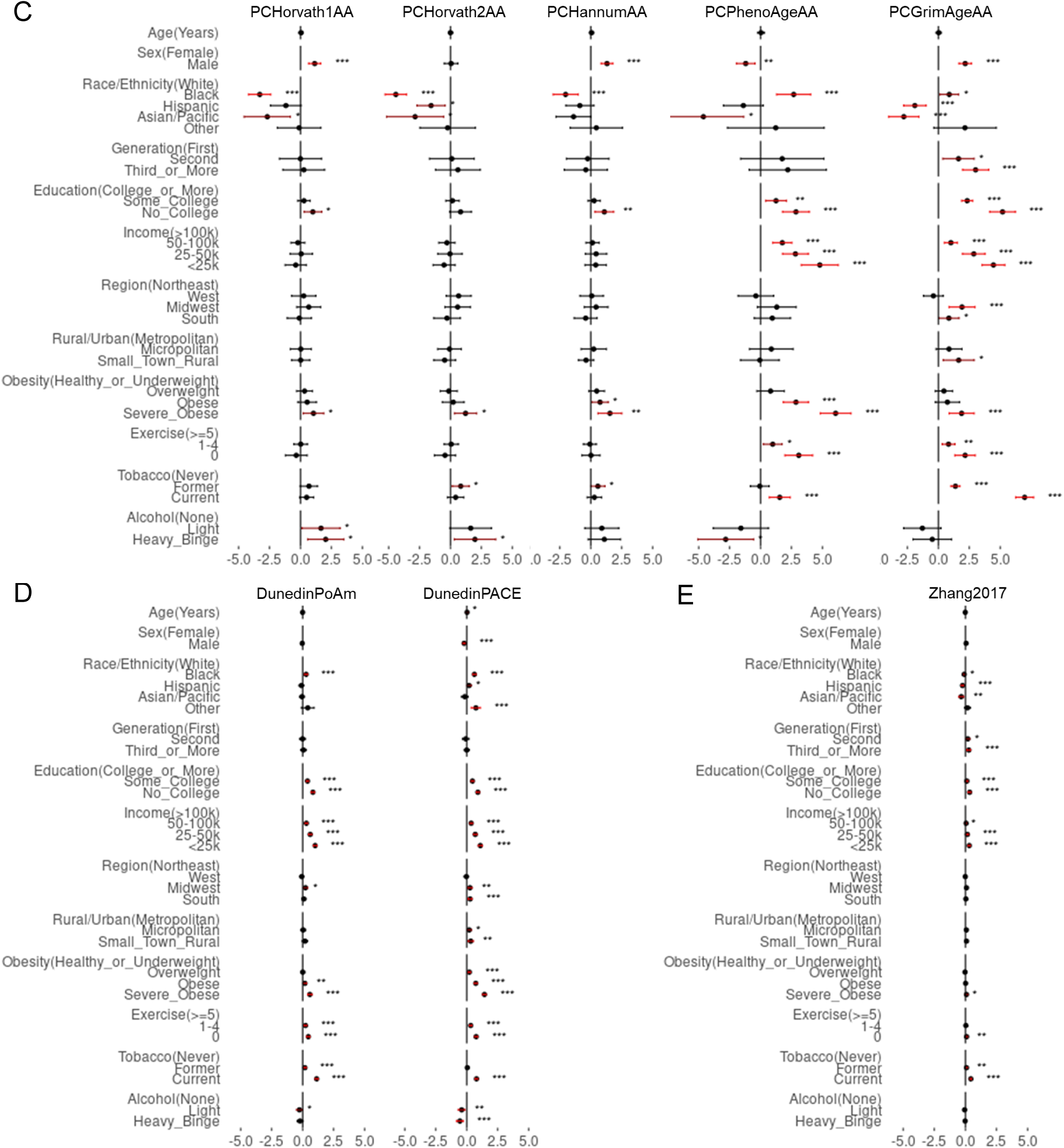
Predicted epigenetic clock measures from weighted bivariate models with socioeconomic and lifestyle factors (N=4237). Notes: First-generation (A), second-generation (B), and PC (C) clock estimates are in years. Third-generation (D) clock estimates (DunedinPoAm, DunedinPACE) are in standard deviation units. Zhang-2017 (E) is in age-adjusted mortality risk units. Significant associations are shown in red point estimates and error bars; estimates to the left of 0.0 indicate slower epigenetic aging and to the right of 0.0 faster epigenetic aging in years relative to chronological age for the category on the sociodemographic or life-style factor compared to the reference category. * p<.01; ** p<.01; *** p<.001.

Overall, males had more rapid biological aging than females across most clocks. Consistent with the literature, there were mixed results for race/ethnicity; seven of the mainly first-generation clocks showed slower biological aging and six of the second- and third-generation clocks showed faster epigenetic aging among Black adults compared to White adults. More consistent results were found for Asian/Pacific Islanders with slower biological aging in 10 clocks, and Hispanics with slower biological aging in 5 clocks compared to Whites. Foreign-born first-generation immigrants also had slower biological aging compared to US-born adults.

For the most part, second- and third-generation clocks showed expected associations where lower levels of education and income were associated with accelerated biological aging. Only a few clocks were significantly associated with geographic locations with accelerated biological aging in the South and/or Midwest compared to the Northeast and in more rural locales compared to urban areas.

The strongest and most consistent lifestyle association with EAA was obesity status. Across 15 clocks, those with severe obesity (e.g., BMI≥40) experienced faster biological aging; 9 of the clocks found faster biological aging among the obese compared to those of normal or underweight status. Weekly exercise was the next most consistent and frequently significant lifestyle association, showing those who got less exercise had accelerated biological aging for 9 clocks. Alcohol and tobacco use results were mixed with some clocks estimating slower aging and some faster aging with greater use. Among the second-generation clocks (except for Lin), however, current or former smokers had accelerated biological aging.

When we tested for independent associations in a multivariable model adjusting for all sociodemographic and lifestyle characteristics (**Table 3**), we found factors were most often significantly associated with second- and third-generation measures of biological aging. Focusing on these clocks and highlighting GrimAgeAA as an example, consistent independent associations showing faster biological aging were found for the following characteristics: those with lower levels of education (no college *β*=2.63, 95%CI (1.67-3.58); some college *β*=.93, 95%CI (0.45-1.40)), and income (<$25,000 *β*=1.70, 95%CI (0.68-2.72); $25,000-$50,000 *β*=1.14, 95%CI (0.18-2.10)); severe obesity status (*β*=1.57, 95%CI (0.51-2.63)); lack of exercise (no bouts/wk *β*=1.33, 95%CI (0.67-1.99); and tobacco use (current smoker *β*=7.16, 95%CI (6.25-8.07)). Most clocks (except PCPhenoAgeAA, DunedinPACE) showed males with faster biological aging (e.g., GrimAgeAA *β*=1.78, 95%CI (1.26-2.30)). Adjusted results for race/ethnicity remained mixed as Black, Asian/Pacific Islander and Hispanic adults had slower rates of biological aging compared to Whites adults across most clocks. The slower biological aging among foreign-born immigrants was independently significant only for the GrimAge clocks, while HannumAA showed slower aging among U.S.-born in the third generation. Models that control for cell composition (**Supplement Table 2**) showed similar results.

**Table 3.** Epigenetic clock estimates from weighted multivariate models of sociodemographic and lifestyle characteristics (N=4237).

| Variable |  | Horvath1AA |  | Horvath2AA |  | HannumAA |  | Vidal BraloAA |  | Zhang2019AA |  | LinAA |  | PhenoAgeAA |  | GrimAgeAA |  |
| --- | --- | --- | --- | --- | --- | --- | --- | --- | --- | --- | --- | --- | --- | --- | --- | --- | --- |
|  |  | Beta | CI | Beta | CI | Beta | CI | Beta | CI | Beta | CI | Beta | CI | Beta | CI | Beta | CI |
| Age | Years | -0.09 | (-0.25-0.06) | -0.02 | (-0.12-0.08) | 0.04 | (-0.08-0.16) | -0.07 | (-0.18-0.04) | 0.09 | (-0.02-0.21) | 0.06 | (-0.17-0.28) | 0.01 | (-0.20-0.22) | -0.10 | (-0.22-0.03) |
| Sex | Female (Ref) |  |  |  |  |  |  |  |  |  |  |  |  |  |  |  |  |
|  | Male | **2.40** | **(1.80-2.99)** | 0.06 | (-0.40-0.51) | **1.00** | **(0.55-1.46)** | 0.11 | (-0.37-0.60) | 0.14 | (-0.32-0.60) | 0.61 | (-0.13-1.34) | -0.47 | (-1.27-0.34) | **1.78** | **(1.26-2.30)** |
| Race | White (Ref) |  |  |  |  |  |  |  |  |  |  |  |  |  |  |  |  |
|  | Black | -0.36 | (-1.17-0.45) | -0.28 | (-0.84-0.28) | **-2.35** | **(-3.10--1.60)** | **-2.03** | **(-2.79--1.27)** | **-1.77** | **(-2.47--1.06)** | **1.40** | **(0.26-2.53)** | **-1.93** | **(-3.19--0.68)** | **1.16** | **(0.51-1.82)** |
|  | Hispanic | -0.99 | (-2.51-0.52) | -0.84 | (-1.84-0.16) | -0.87 | (-2.21-0.46) | **-1.46** | **(-2.68--0.24)** | -0.97 | (-2.14-0.21) | -0.62 | (-2.53-1.30) | -0.84 | (-2.72-1.04) | **-1.45** | **(-2.43--0.47)** |
|  | Asian/Pacific | -0.18 | (-2.06-1.71) | **-1.46** | **(-2.56--0.36)** | **-2.18** | **(-3.57--0.78)** | **-2.91** | **(-4.50--1.31)** | **-2.95** | **(-4.32--1.57)** | -1.38 | (-3.36-0.61) | **-3.09** | **(-5.90--0.29)** | -0.98 | (-2.65-0.68) |
|  | Other | 0.58 | (-1.68-2.84) | -0.45 | (-2.63-1.72) | -0.68 | (-3.51-2.16) | -0.16 | (-3.26-2.94) | -0.40 | (-2.26-1.45) | 0.87 | (-2.96-4.71) | 1.31 | (-2.22-4.83) | 0.10 | (-1.94-2.14) |
| Generation | First (Ref) |  |  |  |  |  |  |  |  |  |  |  |  |  |  |  |  |
|  | Second | 0.35 | (-1.88-2.57) | -0.26 | (-1.65-1.13) | -0.86 | (-2.18-0.46) | 0.88 | (-0.66-2.42) | 0.43 | (-0.80-1.66) | 0.33 | (-1.77-2.43) | -0.03 | (-2.73-2.66) | **1.66** | **(0.36-2.96)** |
|  | Third+ | 0.38 | (-2.01-2.77) | -0.57 | (-2.05-0.91) | **-2.07** | **(-3.64--0.49)** | 0.79 | (-0.84-2.42) | 0.01 | (-1.32-1.34) | -0.99 | (-3.28-1.29) | -0.84 | (-3.78-2.11) | **1.72** | **(0.38-3.07)** |
| Education | College+ (Ref) |  |  |  |  |  |  |  |  |  |  |  |  |  |  |  |  |
|  | Some_College | -0.12 | (-0.76-0.51) | 0.17 | (-0.29-0.63) | 0.07 | (-0.43-0.57) | 0.27 | (-0.24-0.77) | 0.16 | (-0.28-0.60) | -0.16 | (-0.99-0.67) | -0.05 | (-0.77-0.67) | **0.93** | **(0.45-1.40)** |
|  | No College | 0.44 | (-0.50-1.39) | 0.42 | (-0.23-1.07) | 0.65 | (-0.19-1.49) | -0.12 | (-1.02-0.78) | -0.09 | (-0.91-0.73) | 0.42 | (-0.85-1.69) | 0.83 | (-0.47-2.13) | **2.63** | **(1.67-3.58)** |
| Income | >100k (Ref) |  |  |  |  |  |  |  |  |  |  |  |  |  |  |  |  |
|  | 50-100k | **-0.78** | **(-1.40--0.15)** | -0.04 | (-0.43-0.35) | 0.52 | (-0.06-1.09) | -0.36 | (-0.84-0.13) | -0.08 | (-0.58-0.43) | -0.52 | (-1.69-0.66) | 0.60 | (-0.21-1.41) | 0.28 | (-0.30-0.86) |
|  | 25-50k | -0.21 | (-1.07-0.65) | 0.14 | (-0.36-0.64) | 0.80 | (-0.06-1.67) | -0.25 | (-1.02-0.52) | 0.43 | (-0.29-1.14) | -0.18 | (-1.45-1.09) | 1.13 | (-0.16-2.41) | **1.14** | **(0.18-2.10)** |
|  | <25k | -0.17 | (-1.08-0.74) | -0.09 | (-0.72-0.53) | **1.12** | **(0.41-1.83)** | -0.49 | (-1.45-0.46) | 0.45 | (-0.26-1.16) | -0.88 | (-2.26-0.50) | **1.99** | **(0.45-3.52)** | **1.70** | **(0.68-2.72)** |
| Region | Northeast (Ref) |  |  |  |  |  |  |  |  |  |  |  |  |  |  |  |  |
|  | West | 0.63 | (-0.27-1.53) | 0.08 | (-0.51-0.67) | 0.18 | (-0.47-0.84) | -0.40 | (-1.09-0.29) | 0.06 | (-0.60-0.71) | -0.57 | (-1.56-0.42) | 0.90 | (-0.34-2.14) | 0.22 | (-0.56-0.99) |
|  | Midwest | 0.80 | (-0.07-1.68) | -0.05 | (-0.60-0.50) | 0.35 | (-0.27-0.96) | -0.67 | (-1.36-0.01) | 0.23 | (-0.42-0.88) | 0.86 | (-0.27-1.99) | **2.08** | **(0.88-3.29)** | **0.95** | **(0.23-1.67)** |
|  | South | 0.18 | (-0.66-1.02) | 0.34 | (-0.22-0.89) | 0.22 | (-0.45-0.89) | -0.18 | (-0.81-0.45) | 0.35 | (-0.23-0.92) | -0.71 | (-1.64-0.22) | 0.66 | (-0.48-1.79) | 0.00 | (-0.69-0.70) |
| Rural/Urban | Metropolitan (Ref) |  |  |  |  |  |  |  |  |  |  |  |  |  |  |  |  |
|  | Micropolitan | 0.37 | (-0.67-1.41) | 0.42 | (-0.31-1.16) | 0.50 | (-0.42-1.42) | 0.47 | (-0.17-1.11) | 0.31 | (-0.38-1.00) | 0.53 | (-1.21-2.27) | 0.29 | (-1.05-1.62) | 0.46 | (-0.43-1.36) |
|  | Small town/Rural | **-1.07** | **(-1.97--0.17)** | -0.34 | (-0.94-0.25) | -0.56 | (-1.26-0.13) | -0.61 | (-2.04-0.83) | -0.57 | (-1.28-0.14) | -0.69 | (-1.78-0.41) | -0.59 | (-2.08-0.91) | 0.26 | (-0.60-1.12) |
| Obesity | Normal or underweight (Ref) |  |  |  |  |  |  |  |  |  |  |  |  |  |  |  |  |
|  | Overweight | -0.08 | (-0.85-0.70) | 0.32 | (-0.24-0.88) | 0.15 | (-0.37-0.66) | 0.38 | (-0.22-0.97) | 0.14 | (-0.44-0.71) | 0.61 | (-0.37-1.60) | 0.81 | (-0.13-1.75) | 0.11 | (-0.53-0.75) |
|  | Obese | 0.65 | (-0.11-1.42) | **0.87** | **(0.22-1.52)** | **0.82** | **(0.15-1.50)** | **1.08** | **(0.51-1.66)** | 0.66 | (0.00-1.33) | 0.29 | (-0.68-1.27) | **2.13** | **(0.98-3.28)** | 0.05 | (-1.01-1.10) |
|  | Severe Obesity | **2.93** | **(1.69-4.18)** | **0.90** | **(0.11-1.69)** | **2.14** | **(1.14-3.14)** | **1.92** | **(0.98-2.86)** | 0.60 | (-0.21-1.41) | **2.64** | **(1.16-4.12)** | **5.24** | **(3.55-6.92)** | **1.57** | **(0.51-2.63)** |
| Exercise | 5 or more bouts/week (Ref) |  |  |  |  |  |  |  |  |  |  |  |  |  |  |  |  |
|  | 1-4 bouts/week | -0.05 | (-0.66-0.56) | -0.17 | (-0.56-0.22) | 0.29 | (-0.14-0.73) | -0.18 | (-0.67-0.32) | 0.22 | (-0.26-0.69) | 0.02 | (-0.78-0.82) | 0.55 | (-0.22-1.31) | **0.52** | **(0.03-1.02)** |
|  | No bouts/week | 0.29 | (-0.35-0.92) | 0.24 | (-0.34-0.82) | 0.58 | (-0.05-1.21) | -0.10 | (-0.75-0.54) | 0.38 | (-0.27-1.03) | **0.98** | **(0.02-1.94)** | **1.84** | **(0.65-3.04)** | **1.33** | **(0.67-1.99)** |
| Tobacco | Never (Ref) |  |  |  |  |  |  |  |  |  |  |  |  |  |  |  |  |
|  | Former | -0.46 | (-1.27-0.35) | **-0.74** | **(-1.33--0.14)** | -0.17 | (-0.68-0.33) | 0.05 | (-0.55-0.64) | **-0.65** | **(-1.20--0.10)** | -0.78 | (-2.01-0.44) | -0.15 | (-0.92-0.61) | **1.57** | **(1.13-2.01)** |
|  | Current | -0.54 | (-1.33-0.25) | **-0.67** | **(-1.25--0.09)** | -0.06 | (-0.85-0.72) | **-0.69** | **(-1.33--0.05)** | **-1.34** | **(-2.04--0.64)** | **-1.88** | **(-2.98--0.78)** | **1.38** | **(0.32-2.44)** | **7.16** | **(6.25-8.07)** |
| Alcohol | None (Ref) |  |  |  |  |  |  |  |  |  |  |  |  |  |  |  |  |
|  | Light | 0.61 | (-1.46-2.68) | 0.83 | (-0.62-2.28) | 0.93 | (-0.18-2.04) | -0.62 | (-2.23-0.98) | 1.06 | (-0.14-2.27) | 0.57 | (-2.02-3.16) | -0.81 | (-3.43-1.82) | -0.70 | (-2.01-0.60) |
|  | Heavy/Binge | 0.51 | (-1.47-2.50) | 0.62 | (-0.87-2.11) | 0.65 | (-0.48-1.76) | -0.82 | (-2.41-0.76) | 1.01 | (-0.20-2.22) | 0.19 | (-2.39-2.78) | -0.79 | (-3.34-1.76) | -0.80 | (-2.12-0.52) |

**Table 3 (Cont.). Epigenetic clock estimates from weighted multivariate models of sociodemographic and lifestyle characteristics (N=4237).**
| Variable |  | PCHorvath1AA |  | PCHorvath2AA |  | PCHannumAA |  | PCPhenoAgeAA |  | PCGrimAgeAA |  | DunedinPoAm |  | DunedinPACE |  | Zhang2017 |  |
| --- | --- | --- | --- | --- | --- | --- | --- | --- | --- | --- | --- | --- | --- | --- | --- | --- | --- |
|  |  | Beta | CI | Beta | CI | Beta | CI | Beta | CI | Beta | CI | Beta | CI | Beta | CI | Beta | CI |
| Age | Years | 0.11 | (-0.03-0.24) | 0.09 | (-0.04-0.23) | 0.09 | (-0.05-0.23) | 0.00 | (-0.21-0.21) | -0.02 | (-0.15-0.11) | -0.01 | (-0.03-0.02) | 0.02 | (0.00-0.04) | -0.01 | (-0.02-0.01) |
| Sex | Female (Ref) |  |  |  |  |  |  |  |  |  |  |  |  |  |  |  |  |
|  | Male | **1.14** | **(0.60-1.68)** | 0.07 | (-0.50-0.64) | **1.34** | **(0.83-1.85)** | **-0.96** | **(-1.70--0.21)** | **1.90** | **(1.48-2.32)** | -0.06 | (-0.14-0.01) | **-0.23** | **(-0.29--0.16)** | 0.05 | (-0.01-0.10) |
| Race | White (Ref) |  |  |  |  |  |  |  |  |  |  |  |  |  |  |  |  |
|  | Black | **-3.54** | **(-4.32--2.76)** | **-4.68** | **(-5.51--3.84)** | **-2.37** | **(-3.26--1.47)** | **1.33** | **(0.16-2.50)** | 0.30 | (-0.29-0.89) | **0.12** | **(0.01-0.24)** | **0.33** | **(0.22-0.44)** | **-0.16** | **(-0.23--0.08)** |
|  | Hispanic | **-1.68** | **(-2.98--0.38)** | **-1.94** | **(-3.25--0.64)** | **-1.68** | **(-2.89--0.47)** | **-1.98** | **(-3.60--0.35)** | -0.90 | (-1.82-0.02) | -0.07 | (-0.26-0.11) | **0.20** | **(0.04-0.36)** | **-0.19** | **(-0.30--0.09)** |
|  | Asian/Pacific | **-2.89** | **(-4.60--1.18)** | **-2.99** | **(-5.06--0.92)** | **-2.01** | **(-3.50--0.53)** | **-4.23** | **(-7.25--1.21)** | -0.61 | (-1.81-0.58) | 0.13 | (-0.10-0.36) | 0.14 | (-0.13-0.41) | **-0.22** | **(-0.40--0.04)** |
|  | Other | -0.69 | (-2.48-1.09) | -0.59 | (-2.82-1.63) | -0.24 | (-2.32-1.84) | 0.95 | (-2.67-4.57) | 0.50 | (-1.31-2.30) | 0.19 | (-0.18-0.55) | **0.60** | **(0.35-0.86)** | 0.06 | (-0.14-0.26) |
| Generation | First (Ref) |  |  |  |  |  |  |  |  |  |  |  |  |  |  |  |  |
|  | Second | -0.46 | (-1.89-0.98) | -0.43 | (-1.90-1.04) | -0.51 | (-1.93-0.90) | 0.81 | (-1.88-3.50) | **1.27** | **(0.18-2.37)** | -0.05 | (-0.27-0.17) | -0.07 | (-0.29-0.15) | 0.13 | (-0.02-0.27) |
|  | Third+ | -0.89 | (-2.31-0.53) | -0.55 | (-2.08-0.97) | -1.38 | (-2.96-0.20) | -0.33 | (-2.93-2.27) | **1.63** | **(0.62-2.64)** | -0.11 | (-0.33-0.11) | -0.06 | (-0.28-0.16) | 0.10 | (-0.04-0.25) |
| Education | College+ (Ref) |  |  |  |  |  |  |  |  |  |  |  |  |  |  |  |  |
|  | Some College | 0.25 | (-0.22-0.73) | 0.22 | (-0.34-0.78) | 0.09 | (-0.41-0.59) | -0.07 | (-0.91-0.77) | **0.76** | (0.30-1.22) | 0.07 | (-0.02-0.15) | **0.11** | **(0.02-0.19)** | 0.03 | (-0.03-0.10) |
|  | No College | 0.82 | (-0.10-1.75) | 0.83 | (-0.31-1.96) | 0.69 | (-0.28-1.66) | 0.77 | (-0.71-2.25) | **2.10** | (1.20-3.00) | **0.24** | **(0.10-0.37)** | **0.30** | **(0.18-0.42)** | **0.15** | **(0.04-0.27)** |
| Income | >100k (Ref) |  |  |  |  |  |  |  |  |  |  |  |  |  |  |  |  |
|  | 50-100k | -0.15 | (-0.69-0.39) | -0.17 | (-0.79-0.45) | 0.18 | (-0.37-0.72) | **0.82** | **(0.07-1.57)** | 0.16 | (-0.25-0.58) | **0.14** | **(0.06-0.23)** | 0.07 | (-0.01-0.15) | 0.02 | (-0.04-0.07) |
|  | 25-50k | 0.35 | (-0.62-1.32) | 0.45 | (-0.64-1.54) | 0.63 | (-0.29-1.54) | **1.29** | **(0.15-2.42)** | **1.01** | **(0.17-1.85)** | **0.31** | **(0.19-0.43)** | **0.17** | **(0.06-0.28)** | 0.08 | (-0.02-0.18) |
|  | <25k | 0.38 | (-0.72-1.47) | 0.47 | (-0.68-1.61) | 0.94 | (-0.02-1.90) | **2.47** | **(0.88-4.05)** | **1.29** | **(0.39-2.19)** | **0.48** | **(0.33-0.63)** | **0.33** | **(0.17-0.48)** | **0.16** | **(0.05-0.27)** |
| Region | Northeast (Ref) |  |  |  |  |  |  |  |  |  |  |  |  |  |  |  |  |
|  | West | 0.35 | (-0.51-1.20) | 0.76 | (-0.15-1.67) | 0.13 | (-0.74-1.00) | 0.16 | (-1.05-1.37) | -0.13 | (-0.73-0.47) | -0.01 | (-0.13-0.12) | 0.06 | (-0.04-0.16) | 0.02 | (-0.08-0.11) |
|  | Midwest | 0.44 | (-0.37-1.24) | 0.31 | (-0.60-1.22) | 0.28 | (-0.54-1.11) | 0.79 | (-0.32-1.89) | **0.69** | **(0.09-1.28)** | 0.08 | (-0.04-0.21) | **0.13** | **(0.05-0.22)** | 0.00 | (-0.09-0.09) |
|  | South | 0.32 | (-0.47-1.10) | 0.36 | (-0.54-1.26) | -0.19 | (-1.00-0.62) | 0.20 | (-0.88-1.29) | -0.13 | (-0.64-0.39) | -0.01 | (-0.13-0.10) | **0.10** | **(0.01-0.19)** | 0.00 | (-0.09-0.08) |
| Rural/Urban | Metropolitan (Ref) |  |  |  |  |  |  |  |  |  |  |  |  |  |  |  |  |
|  | Micropolitan | 0.25 | (-0.58-1.09) | 0.28 | (-0.66-1.22) | 0.35 | (-0.68-1.39) | 0.06 | (-1.34-1.46) | 0.02 | (-0.75-0.79) | -0.10 | (-0.26-0.06) | 0.03 | (-0.07-0.13) | 0.03 | (-0.05-0.12) |
|  | Small town/Rural | -0.26 | (-0.94-0.43) | -0.79 | (-1.62-0.05) | **-0.65** | **(-1.27--0.03)** | -1.31 | (-2.80-0.17) | 0.19 | (-0.57-0.95) | -0.06 | (-0.22-0.10) | 0.07 | (-0.09-0.23) | -0.02 | (-0.13-0.09) |
| Obesity | Normal or underweight (Ref) |  |  |  |  |  |  |  |  |  |  |  |  |  |  |  |  |
|  | Overweight | 0.29 | (-0.37-0.96) | 0.12 | (-0.54-0.79) | 0.37 | (-0.26-0.99) | 0.88 | (-0.10-1.85) | 0.01 | (-0.49-0.50) | 0.01 | (-0.09-0.12) | **0.21** | **(0.12-0.30)** | -0.03 | (-0.10-0.04) |
|  | Obese | 0.66 | (-0.06-1.39) | 0.62 | (-0.23-1.48) | **0.72** | **(0.04-1.40)** | **2.19** | **(1.25-3.14)** | -0.22 | (-1.01-0.58) | 0.05 | (-0.05-0.15) | **0.58** | **(0.48-0.68)** | -0.01 | (-0.10-0.07) |
|  | Severe Obesity | **1.47** | **(0.66-2.29)** | **1.85** | **(0.97-2.73)** | **1.74** | **(0.80-2.69)** | **4.73** | **(3.46-5.99)** | 0.86 | (0.00-1.73) | **0.39** | **(0.24-0.54)** | **1.16** | **(1.04-1.28)** | 0.05 | (-0.04-0.14) |
| Exercise | 5 or more bouts/week (Ref) |  |  |  |  |  |  |  |  |  |  |  |  |  |  |  |  |
|  | 1-4 bouts/week | -0.03 | (-0.51-0.46) | -0.04 | (-0.55-0.46) | -0.15 | (-0.63-0.33) | 0.24 | (-0.46-0.93) | 0.34 | (-0.09-0.78) | 0.08 | (0.00-0.16) | **0.11** | **(0.03-0.18)** | 0.01 | (-0.05-0.07) |
|  | No bouts/week | 0.09 | (-0.69-0.86) | -0.05 | (-0.80-0.70) | 0.19 | (-0.47-0.86) | **1.22** | **(0.20-2.23)** | **1.19** | **(0.56-1.83)** | **0.16** | **(0.05-0.26)** | **0.28** | **(0.18-0.38)** | 0.04 | (-0.03-0.11) |
| Tobacco | Never (Ref) |  |  |  |  |  |  |  |  |  |  |  |  |  |  |  |  |
|  | Former | 0.05 | (-0.60-0.69) | 0.12 | (-0.51-0.74) | 0.16 | (-0.38-0.69) | 0.20 | (-0.56-0.97) | **1.06** | **(0.69-1.44)** | **0.20** | **(0.11-0.29)** | **0.13** | **(0.05-0.20)** | **0.06** | **(0.01-0.11)** |
|  | Current | -0.15 | (-0.78-0.48) | -0.12 | (-0.81-0.57) | -0.33 | (-1.00-0.34) | 0.85 | (-0.11-1.80) | **5.64** | **(4.90-6.38)** | **0.96** | **(0.84-1.08)** | **0.66** | **(0.55-0.77)** | **0.34** | **(0.28-0.41)** |
| Alcohol | None (Ref) |  |  |  |  |  |  |  |  |  |  |  |  |  |  |  |  |
|  | Light | **1.62** | **(0.01-3.23)** | 1.26 | (-0.21-2.73) | 1.12 | (-0.35-2.59) | -0.46 | (-2.79-1.86) | -0.37 | (-1.62-0.87) | -0.08 | (-0.26-0.10) | -0.12 | (-0.32-0.09) | -0.02 | (-0.15-0.11) |
|  | Heavy/Binge | **1.83** | **(0.26-3.39)** | 1.52 | (0.00-3.04) | 1.26 | (-0.18-2.70) | -1.14 | (-3.46-1.19) | -0.25 | (-1.43-0.92) | -0.05 | (-0.24-0.14) | -0.19 | (-0.40-0.02) | -0.02 | (-0.14-0.11) |

Notes: Controls for cell composition not included. Bolded estimates are significant (at p<.05 or less).

## Discussion

Based on the current literature, we estimated 16 epigenetic clocks, primarily developed for older aged cohorts, to assess their distribution, correlations with chronological age and with each other, and their variability across sociodemographic and lifestyle characteristics known to predict morbidity and mortality in prior research using the younger adult Add Health cohort. While it makes sense that most epigenetic research focuses on older adults, there is growing recognition that molecular processes underlying disease risk begin long before overt disease is evident in chronologically older aged adults.^52,53^

Our results found that the clock measures displayed a range of predicted epigenetic ages for the younger adults that were moderately correlated with chronological age, and with each other, and this result is consistent with prior epigenetic clock research on older samples.^9,11^ These wide-ranging results suggest that different clocks may reflect distinct aspects of aging given they are based on the assessment of methylation at highly disparate numbers of CpG sites, trained on different populations that vary by age, race and ethnicity, and developed in different tissues.^5^

Nevertheless, many social and lifestyle factors were associated with second- and third-generation clocks in the expected direction according to prior research on inequalities in health and mortality risks,^26,54,55^ even in this sample of adults about to enter midlife. In particular, GrimAge, PCGrimAge, and DunedinPACE showed accelerated aging among those with no or some college education relative to a college degree and for those at near or below poverty-level incomes relative to those with incomes above $100,000. Importantly and consistent with other research, severe obesity and lack of weekly exercise were also associated with faster biological aging for second- and third-generation clocks. Given second- and third-generation clocks were trained to predict disease and mortality risks as opposed to first-generation clocks that were trained on chronological age, our findings support the conclusion that the biological aging process is underway prior to mid- and later life and these second- and third-generation clocks are sensitive measures of this process before age-related disease comorbidities are present.

We found interesting new results for immigrant status. Despite the often stressful and discriminatory contexts in which immigrants live, US-born status, even those with foreign-born parents in the second generation, appear to experience faster aging, suggesting the immigrant advantage found in much prior research remains biologically embedded.^56,57^ Note, however, this result may be driven by the large and heterogenous Hispanic population that comprised the immigrant groups as Hispanics tend to have lower biological aging for second- and third-generation clocks.

One potential limitation of our research is the small age range of adults in Add Health. This limitation could be related to inconsistent results we find for race, ethnicity, and sex (though this finding is also consistent with prior research^5,9,11,35,58,59^). There are differing views on whether epigenetic markers can be analyzed in pooled racial and ethnic samples, especially because many of the epigenetic clocks have been developed in predominantly White samples.^7,13,20,60^ It remains to be investigated whether there are varying responses to social and environmental exposures in different racial and ethnic groups.^60^ Prior research identified changes in biological aging by sex that is related to age (e.g., females experience more rapid aging during menopause^38^), which should not affect our young adult sample. However, females also have variability in immunity over time—especially during and after pregnancy - and experience variation in autoimmunity and hormones,^61^ which could be related to some of the outcomes on which the clocks are or are not trained. In addition, other social factors we did not include may explain some of the mixed sex and race/ethnicity findings.^60^

There remains great promise in research using epigenetic clocks to understand the underlying molecular and cellular changes that accompany aging processes and the development of chronic disease. Future epigenetic research should prioritize representative and diverse samples such as Add Health to evaluate whether cellular and molecular changes do indeed vary by sex, race, and ethnicity so we can improve the measurement of biological aging. Our study further showed cellular change in underlying disease processes was already evident in younger adults which can inform prevention efforts. Future research should therefore include a broader range of ages to allow investigation of potential moderation by age or life stage to better understand when and how social inequalities in biological aging emerge across the life course.

## Acknowledgements

This project was funded by the National Institutes of Health (NIH) National Institute on Minority Health and Health Disparities R01MD013349 (Harris and Aiello) and by the *Eunice Kennedy Shriver* National Institute of Child Health & Human Development F32HD103400 (Mishra). We are also grateful to the Carolina Population Center for general support (P2CHD050924). This research uses data from Add Health, funded by grant P01HD31921 (Harris) from the *Eunice Kennedy Shriver* National Institute of Child Health and Human Development (NICHD), with cooperative funding from 23 other federal agencies and foundations. Add Health is currently directed by Robert A. Hummer and funded by the National Institute on Aging cooperative agreements U01 AG071448 (Hummer) and U01AG071450 (Aiello and Hummer) at the University of North Carolina at Chapel Hill. Add Health was designed by J. Richard Udry, Peter S. Bearman, and Kathleen Mullan Harris at the University of North Carolina at Chapel Hill.

## Conflict of Interest

None.

## Supplement 1. Data Quality and Curation of Epigenetic Data in Add Health

### 1. Purpose

Add Health (the National Longitudinal Study of Adolescent to Adult Health) is a nationally representative cohort study of U.S. adolescents in grades 7-12 in 1994 and followed for 25 years across five interview waves^1^. The Add Health epigenetic data were generated in Wave V (2016-18) when the cohort was aged 33-44.

The purpose of this document is to describe the provenance, quality control, and curation of the Add Health epigenetic data set.

### 2. Participant Sampling

Specimens were collected as part of the Wave V biomarker visit. After participation in the Wave V survey, respondents are visited by a field examiner/phlebotomist to collect physical measures, biological specimens, and a medications inventory. Venous blood (10 mL serum + 10 mL EDTA + 3 mL EDTA + 6 mL potassium oxalate / sodium fluoride + PAXgene sample), was collected via conventional phlebotomy, promptly centrifuged in the field, and securely shipped to the Laboratory for Clinical Biochemistry Research (LCBR) in Vermont for re-centrifugation, aliquoting, biomarker assay, and archival storage.^2^ The methylation subsample included 4,582 young adults with diverse social, biological, environmental, and behavioral longitudinal data from birth, GWAS, transcriptome, and biomarker data.

### 3. Bisulfite arrays

DNA was extracted from blood samples stored at the LCBR laboratory and quantified and quality checked using the PicoGreen dsDNA kit from Thermofisher and the Synergy4 fluorometer before being sent to the Human Genetics Center Core Laboratory at the University of Texas Health Science Center at the University of Texas, Houston (UT Houston) for methylation analysis using the Illumina Infinium chemistry. The quality of DNA was determined by gel electrophoresis and 500 nanograms of the DNA were subjected to bisulfite conversion using the EZ-96 DNAm Kit (Zymo Research Corporation; Irvine, CA, USA). DNAm levels across ∼850,000 sites were measured using the Infinium MethylationEPIC BeadChip (Illumina, Inc.; San Diego, CA). Each plate included control samples, including one positive control (Universal Methylated Human DNA Standard; Zymo Research Corporation; Irvine, CA, USA), one negative control, which is human DNA that has been whole genome amplified with phi29 DNA polymerase to create an unmethylated control, and replicates that allow for evaluation of the consistency of DNAm measurements at individual CpG sites. The resulting data from the chip was read into idat files that indicate the green and red wavelength fluorescence intensity at each site on the EPIC chip and transmitted as data matrices from the UT Houston laboratory to Add Health personnel.

### 4. Post data collection analysis

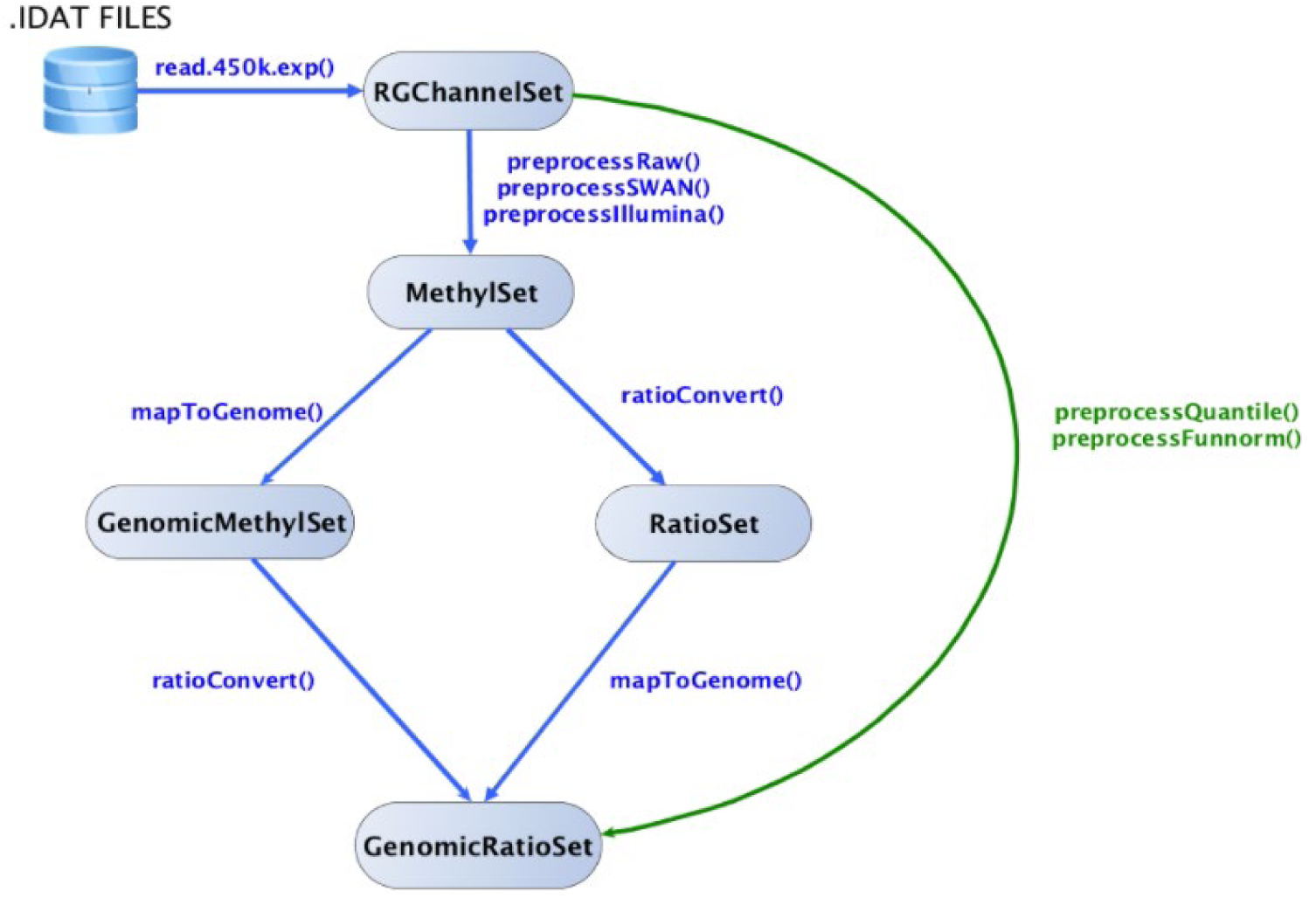

This cartoon shows the interconversions and functions that transform the various data sets that are described here.

#### 4.1 RGChannelSet

The Minfi R bioconductor package was used to generate summarized experiment objects from the red and green fluorescence intensity matrices provided by the laboratory. The first data transformation included the creation of an RGChannelSet object using the following R code:

~~~
library(minfi)
library(IlluminaHumanMethylationEPICmanifest)
library(IlluminaHumanMethylationEPICanno.ilm10b4.hg19)
load(’∼/epigenetics1/fromUTH/EPIC.methylumi.minfi.RData’)
green = rawSet@assays@data@listData$Green
red = rawSet@assays@data@listData$Red
annotation = annotation(rawSet)
rgset=RGChannelSet(Green=green, Red=red, annotation=annotation)
~~~

This object contains the following manifest metadata:

~~~
IlluminaMethylationManifest object
Annotation
 array: IlluminaHumanMethylationEPIC
Number of type I probes: 142262
Number of type II probes: 724574
Number of control probes: 635
Number of SNP type I probes: 21
Number of SNP type II probes: 38
~~~

Additionally, the metadata for this summarized object is as follows:

~~~
class: RGChannelSet
dim: 1051539 2022
metadata(0):
assays(2): Green Red
rownames(1051539): 1600101 1600111 … 99810990 99810992
rowData names(0):
colnames(2022): 204068280089_R03C01 204068280089_R04C01 …
 204074220145_R02C01 204073570043_R04C01
colData names(0):
Annotation
 array: IlluminaHumanMethylationEPIC
 annotation: ilm10b4.hg19
~~~

#### 4.2 Methylset

The rgset object was then converted to a methyset object that contains the unmethylated to methylated signals for each participant at each site. Further, this implements a background subtraction method that estimates background noise from the out-of-band probes and remove it for each sample separately, while the dye-bias normalization utilizes a subset of the control probes to estimate the dye bias. The mset object was generated using the following R code:

~~~
mset = preprocessNoob(rgset)
~~~

The mset object metadata is as follows:

~~~
class: MethylSet
dim: 865859 2022
metadata(0):
assays(2): Meth Unmeth
rownames(865859): cg18478105 cg09835024 … cg10633746 cg12623625
rowData names(0):
colnames(2022): 204068280089_R03C01 204068280089_R04C01 …
 204074220145_R02C01 204073570043_R04C01
colData names(0):
Annotation
 array: IlluminaHumanMethylationEPIC
 annotation: ilm10b4.hg19
Preprocessing
 Method: NA
 minfi version: NA
 Manifest version: NA
~~~

#### 4.3 Genomic RatioSet

The mset object was converted to a genomic ratioset which may be used to extract betas, copy number, and m values. The function preprocessFunnorm implements a functional normalization algorithm that uses the internal control probes to infer between-array technical variation. By default, preprocessFunnorm applies the preprocessNoob function as a first step for background subtraction and uses the first two principal components of the control probes to infer the unwanted variation. This process maps the methylation signals to the genome and is implemented using the following R code:

~~~
grset = preprocessFunnorm(rgset)
~~~

The grset object metadata is as follows:

~~~
class: GenomicRatioSet
dim: 865859 2022
metadata(0):
assays(2): Beta CN
rownames(865859): cg14817997 cg26928153 … cg07587934 cg16855331
rowData names(0):
colnames(2022): 204068280089_R03C01 204068280089_R04C01 …
 204074220145_R02C01 204073570043_R04C01
colData names(3): xMed yMed predictedSex
Annotation
 array: IlluminaHumanMethylationEPIC
 annotation: ilm10b4.hg19
Preprocessing
 Method: NA
 minfi version: NA
 Manifest version: NA
~~~

The betas, copy number (cn), and m values are extracted from the grset object. Some of the CpG sites overlap with short nucleotide polymorphisms and must be censored prior to betas, cn, and m value generation. This is done using the following R code:

~~~
grset = dropLociWithSnps(grset, snps=c(“SBE”,“CpG”), maf=0)
beta = getBeta(grset)
m = getM(grset)
cn = getCN(grset)
~~~

#### 4.4 Quality Control

##### 4.4.1 Intensity plot

The Minfi package contains a standardized set of quality control analysis functions. The first of these functions determines if there exists an imbalance in the fluorescence intensity between methylated and unmethylated sites for any individual in the dataset. This was conducted using the following R code and resulted in the following plot:

~~~
qc = getQC(mset)
plot(qc)
~~~

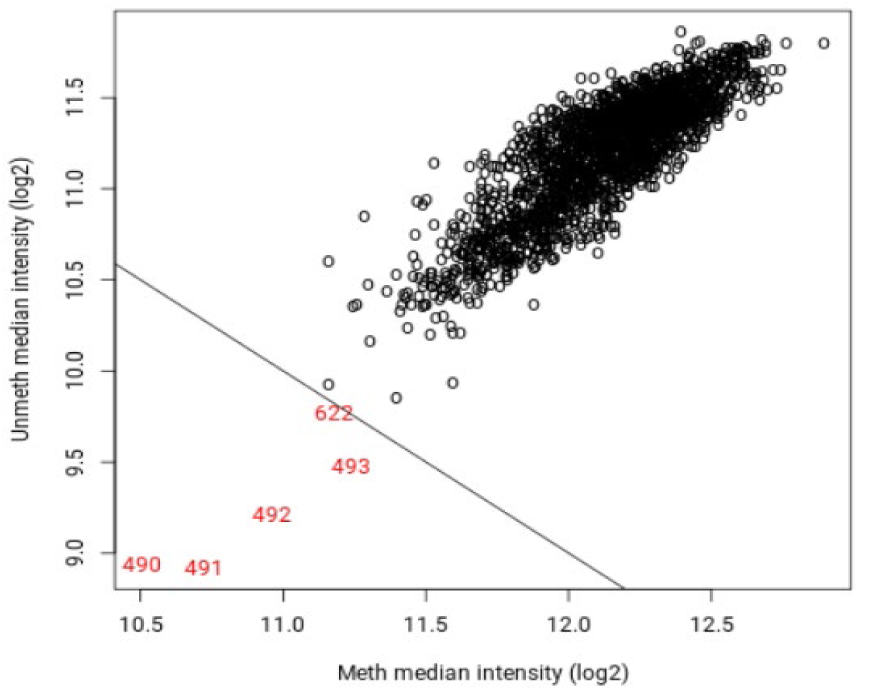

This indicates that there are 5 individuals who must be flagged because their signal intensities were below the expected rates and represent possible outliers.

##### 4.4.2 Density plot

The second Minfi quality control function generates a density plot which analyzes if any of the samples have high levels of hemimethylation, that is, partially methylated sites.

This density plot is produced using the following R code and generates the following plot:

~~~
densityPlot(mset)
~~~

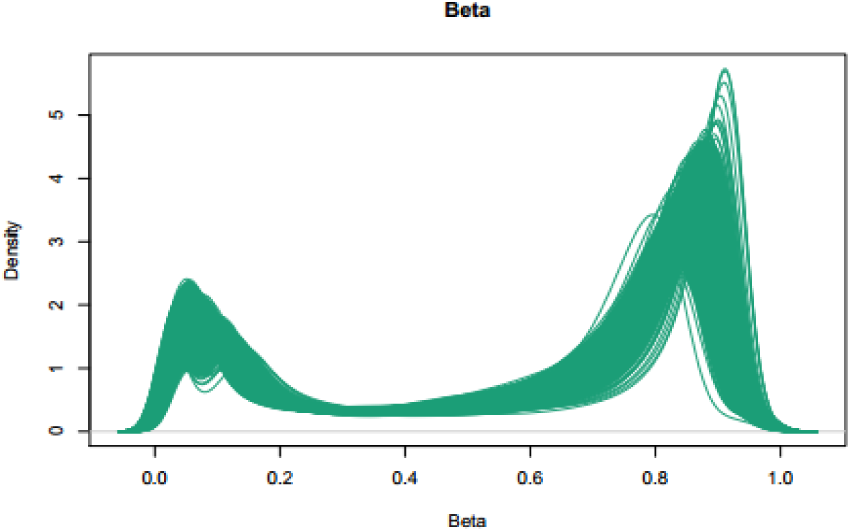

This output indicates there were no samples with an intermediate level of methylation and instead two clear peaks: one for unmethylated signal and one for methylated signal.

##### 4.4.3 Control Strip Plot

This array contains several internal control probes that can be used to assess the quality control of different sample preparation steps (bisulfite conversion, hybridization, etc.). The values of these control probes are stored in the initial RGChannelSet and can be plotted by using the function controlStripPlot and by specifying the control probe type. This is shown for bisulfite conversion using the following R code and generates the following plot:

~~~
controlStripPlot(rgset, controls=“BISULFITE CONVERSION II”)
~~~

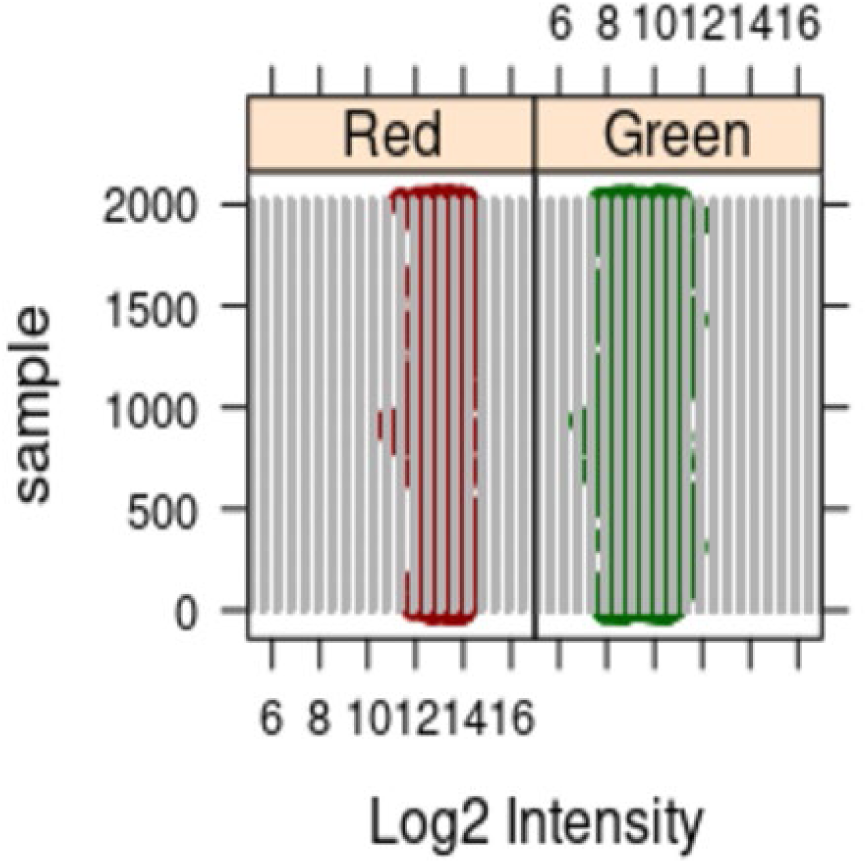

This output demonstrates that there are no outlier samples that have particularly high or low signal intensity for the green and red channels.

##### 4.4.4 Sex check

An analysis of the signal intensity of sites on the X and Y sex chromosomes was completed and compared to recorded survey information about sex at birth. This analysis was completed using the following R code:

~~~
predictedSex = getSex(grset, cutoff = -2)$predictedSex
~~~

6 individuals were inconsistent between recorded survey responses about sex at birth and predictedSex and were censored from further analysis.

##### 4.4.5 Multidimensional scaling

The 50000 most variable CpG sites were selected and a multidimensional analysis was conducted to identify outlier individuals. As expected, the most variable sites were primarily in the sex chromosomes and thus clustered males and females separately.

The resulting plot is as follows:

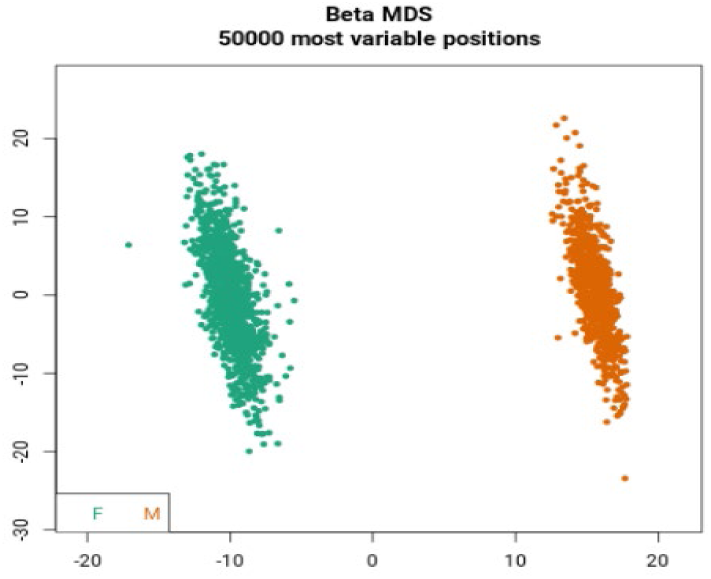

The absence of any obvious outliers and the clear clustering of males and females necessitated no more censoring of individuals.

### 5 Samples and Replicates

#### 5.1 Technical replicates

There were multiple types of embedded controls represented in this data set. The first type of embedded control was technical replicates. These were samples that were analyzed on separate chips but arose from the same sample. There were 51 such samples. The correlation between these samples was analyzed by Spearman rank correlation and presented for batch 1 in the following figure:

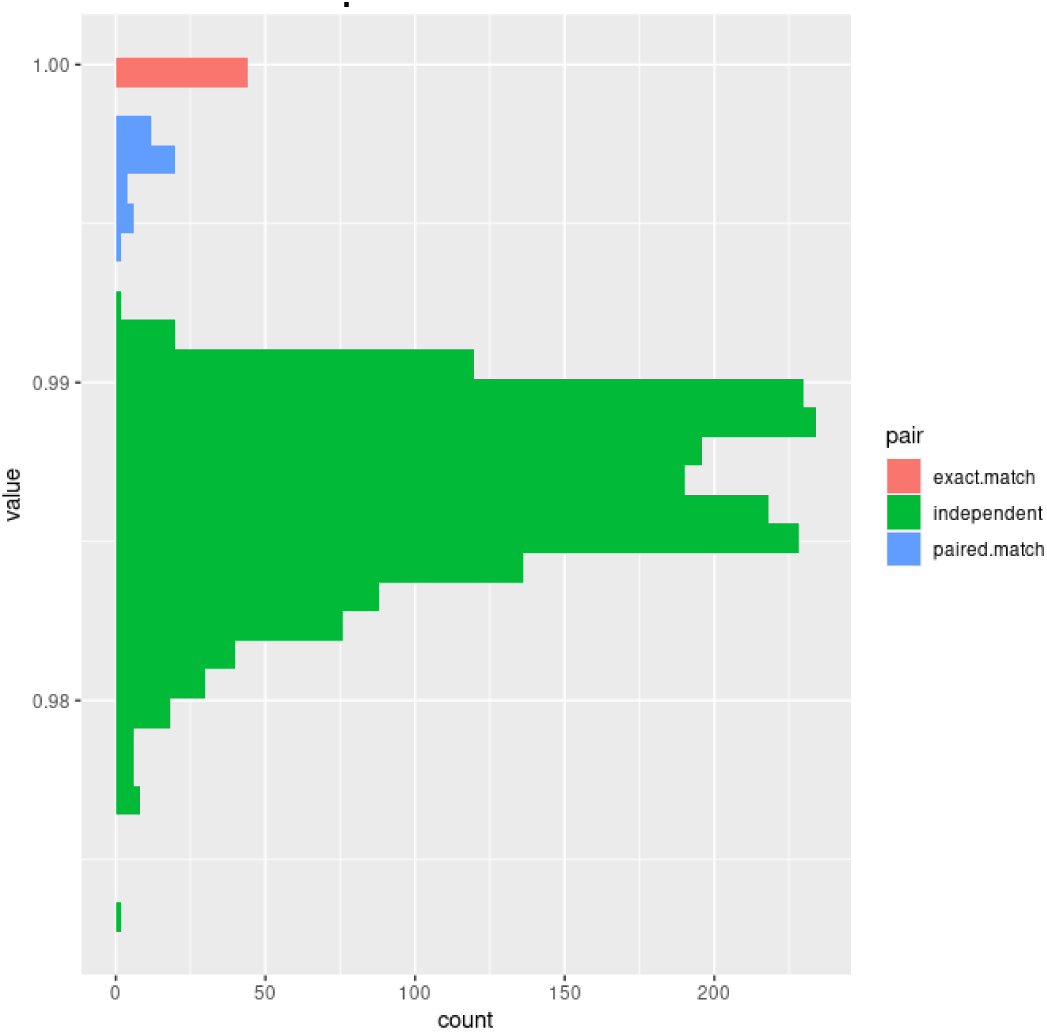

The red indicates samples compared to themselves, which, has a correlation measure of 1.0. The green indicates all comparisons between each of the 51 samples except for paired matches. Finally, blue indicates the correlation of paired samples. While the correlation rates were high for all samples, the correlation between the samples and their paired mate from a different chip were measurably higher and thus provided confidence that the inter-array variability was low.

#### 5.2 Biological replicates

The second type of replicates present in this data set were biological replicates taken from the same participant one week apart. There may be some small changes in methylation patterns at this time scale but they should be swamped by the differences that exist between individuals. There were 200 such samples. The correlation between the first sample and the sample taken one week later as expressed in Spearman rank correlation values for batch1 are as follows:

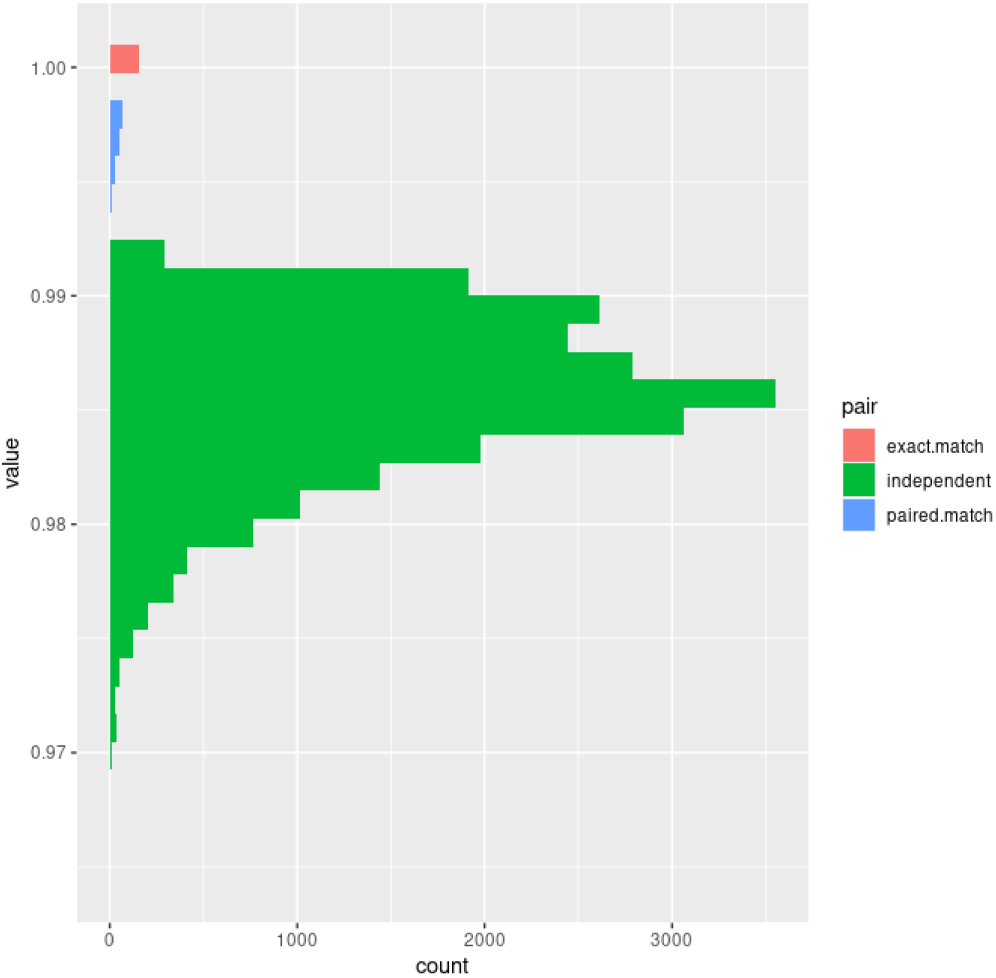

The red indicates samples compared to themselves, with a correlation measure of 1.0. The green indicates all comparisons between independent samples excluding paired matches. Finally, blue indicates the correlation of paired samples. While the correlation rates were high for all samples, the correlation between the samples and their paired mate from one week later were measurably higher and thus provided confidence that the variability in the signal was more related to inter individual differences than technical or chronological variability.

### 6. EPIC 850K Chip

The chip used to generate the data in this dataset is the Illumina EPIC 850K which contains 866,836 epigenetic markers including CpG sites, DNase hypersensitivity regions, SNPs, and various other probe sets. The sites covered by probes on this chip include all autosomes and both sex chromosomes. These sites include most of the sites from other common methylation arrays including the 450K chip as well as 350,000 new probes covering sites annotated by the Fantom5 and ENCODE projects as being important regulatory sites. There are two types of probe sets included in this dataset; type I probes have two separate probe sequences per CpG site (one each for methylated and unmethylated CpGs), whereas type 2 probe sets have just one probe sequence per CpG site. Type 2 probe sets are preferred because of their simplicity but type I probes are retained because they can distinguish methylation in denser regions of CpG sites. Importantly, with the addition of many more probes over previous chips, there is evidence of cross hybridization of some probe sets to related sequences elsewhere in the genome, and these require censoring as described later. Additionally, some probe sets fail to detect their targets at all and must be censored.

### 7. Site Filtering

The initial number of probe sets present in this data set reflect over 850,000 sites in the genome but not all are suitable for use in determining epigenetic age, allostatic load, or differential methylation. Some of these non-suitable sites include SNPs which may include methylation in some participants but not others as a byproduct of their nucleotide sequence. Therefore, all CpG sites overlapping with a SNP were censored from analysis using the following code in R:

~~~
dropLociWithSnps(grset, snps=c(“SBE”,“CpG”), maf=0)
~~~

Further, some of the probes failed to detect signals for enough participants and were also censored. This was achieved using the following code in R:

~~~
detectionP(rgset, type = “m+u”)
~~~

Finally, there is evidence of cross hybridization of some of the probe sets with other sites in the genome. Usage of these probes would complicate interpretation and they were also removed. The 43,000 sites identified as containing cross reactive probes^3^were censored from the final data set.

### 8. Sample filtering

There were 4734 samples initially analyzed for methylation patterns but several samples were removed for quality control purposes. Individual samples were removed for 3 reasons. Any individuals failing the sex check were removed from further analysis, resulting in the censoring of 1 individual. Individuals with low signal intensity from the intensity plot were flagged but not removed (n=5). Lastly, individuals with low sample volumes or concentrations such that the bisulfite conversion was impossible were not assayed. Additionally, technical and biological replicate samples were removed from the final dataset yielding 4582 samples.

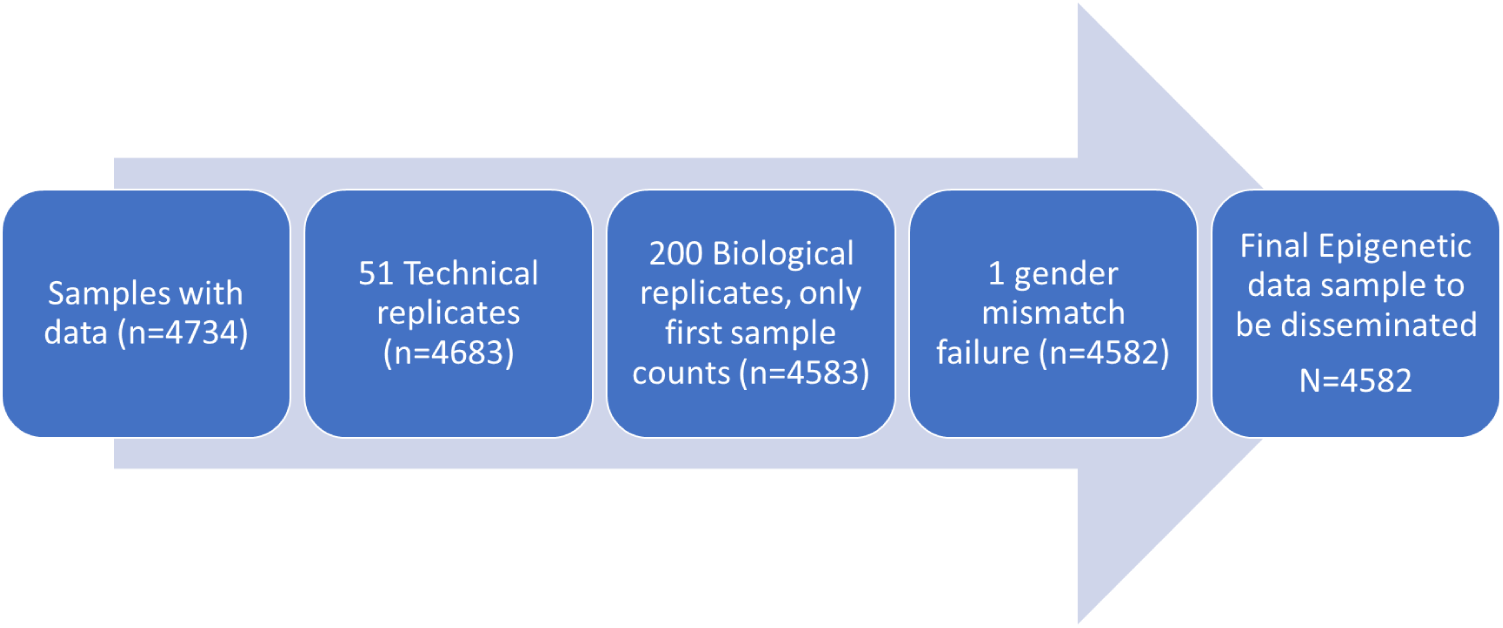

### 9. Batch Correction and Surrogate Variable Adjustments

The R package ComBat was used to correct for batch and the 3 batches were concatenated.

### 10. Epigenetic Age Generation

Background subtracted beta values derived from the ratioset object were used with the Horvath epigenetic age clock to generate an estimated epigenetic age for each participant. The data sets were restricted to a set of 30,484 CpG sites that were relevant for the calculator and the identifier values were dummy coded to maintain data security. Finally, the data was converted to csv with Windows line endings, chunked into groups of 250 individuals, and submitted in batches to the Horvath calculator found at http://dnamage.genetics.ucla.edu/new and used with the methylCIPHER algorithm ^4^ https://github.com/MorganLevineLab/methylCIPHER and Dunedin calculators^5,6^ to generate the epigenetic clocks.

Construction of the PC clocks required an expanded set of 78,464 CpGs, of which 1400 were missing from the processed Add Health betas. Missing CpGs were imputed in R using mean values from GSE40269 ^7^ according to the Levine lab code at https://hub.com/MorganLevineLab/PC-Clocks/, and the PCs and clocks were then constructed using code from the same repository.

### 11. Sample Cell Counts

When a complex tissue such as whole blood is used as a sample, it is important to account for the potential differences in cell types between samples. The Minfi R package was used to calculate the relative amounts of each cell type present in the sample using the following R code:

~~~
cellcounts = estimateCellCounts(rgset, compositeCellType = “Blood”)
~~~

The function analyzes CpG sites whose methylation patterns associate with one of several immune cell subsets including “Bcell”, “CD4T”, “CD8T”, “Granulocytes”, “Monocytes”, and “Natural Killer Cells”. The resulting values for everyone are stored as relative amounts adding up to 1 for everyone.

The resulting values are shown as follows:

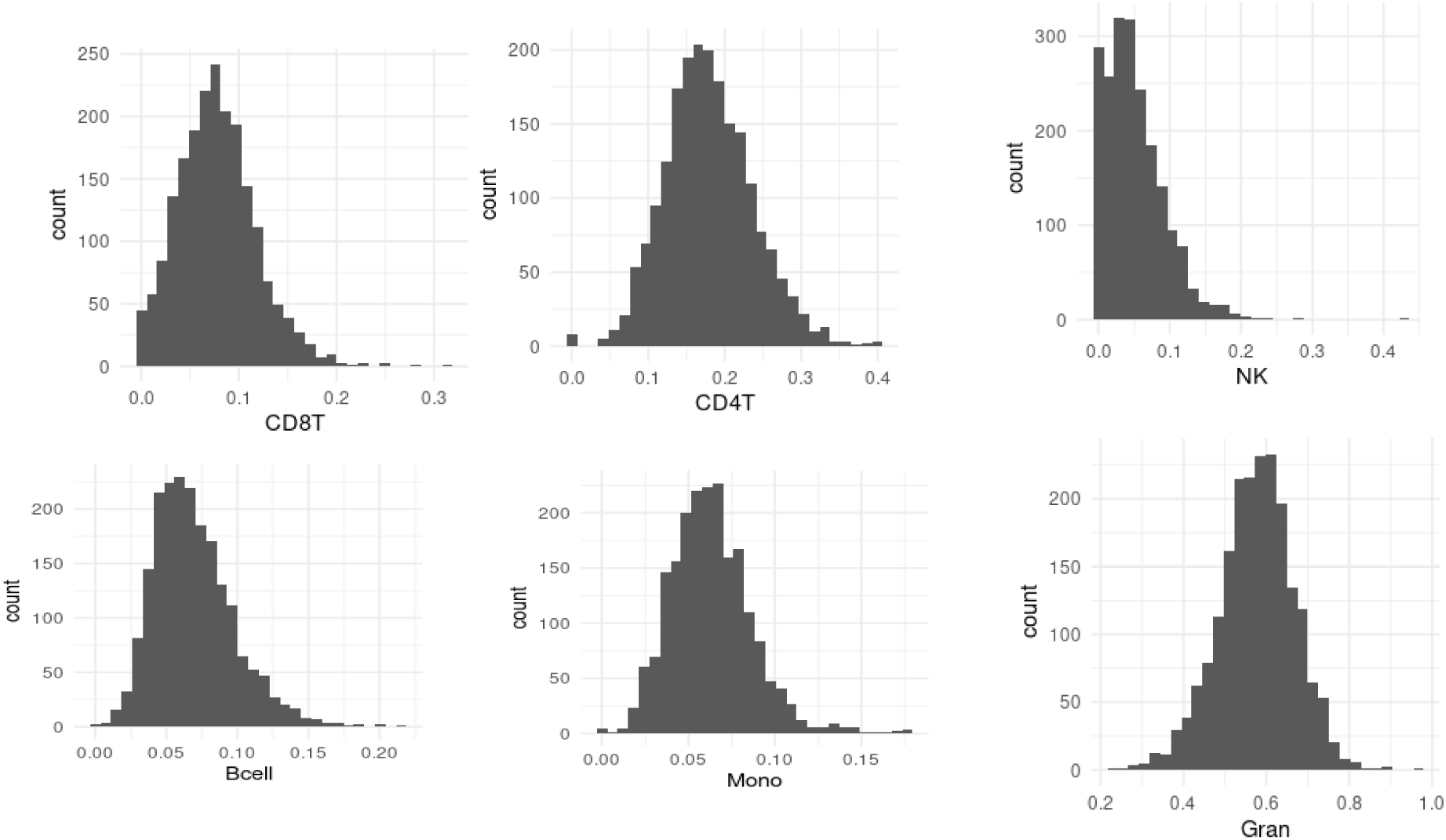

## Supplement 2. Description of Epigenetic Clock and Sociodemographic and Lifestyle Measures

### First Generation Clocks

#### Horvath 1

Horvath 1 is a first-generation epigenetic clock developed in 2013.^1^ This clock was trained in 8,000 samples arising from 82 methylation array datasets, collectively representing 51 healthy tissues and cell types. The clock is calculated from weighted DNA methylation at 353 CpGs present in genes related to cellular survival, proliferation, and tissue development.

Horvath 1 shows strong correlation with chronological age (r=0.96-0.97). This signature is close to 0 for embryonic and pluripotent stem cells and increases with cycles of cellular replication. In 6,000 samples from cancerous tissues, Horvath 1 demonstrated substantial increases in biological age acceleration, averaging several decades older than control samples.

#### Horvath 2

This epigenetic clock, calculated from weighted methylation values at 391 CpGs, better measures the age of human fibroblasts and other skin cells such as keratinocytes, buccal cells, endothelial cells, lymphoblastoid cells, skin, blood, and saliva samples compared to its predecessor.^2^ The improved clock correlates with chronological age in neurons, glia, brain, liver and bone samples. In contrast to the Horvath 1 signature, Horvath 2 predicts biological age acceleration in pathologies related to progeria such as Werner Syndrome. This signature was trained on epigenetic datasets from 10 studies in which the participants had median ages of 0-69 years old.

This clock shares 45 CpGs with the blood-based clock from Hannum^3^ and 60 CpGs with Horvath 1; however, the age acceleration only shows moderate correlations with these two other clocks. Horvath 2 correlates well with cellular passage number but does not show any relationship with telomere length in blood samples.

#### Hannum

Hannum’s epigenetic clock is a blood-based age estimator, calculated from weighted DNA methylation at 71 CpGs.^3^ Hannum et al. developed this clock based on the whole blood of 656 humans at ages 19 to 101 at UCSF, USC, and West China Hospital.

Hannum shows a strong correlation with chronological age (r=0.96) and the rate of DNAm aging is influenced by gender and genetic variants including meQTLs. While this signature was trained on blood samples, it also performs well in breast, kidney, lung and skin samples. Hannum also correlates with gene expression of age-related genes including those involved in developmental biology and DNA repair pathways.

#### VidalBralo

The Vidal-Bralo et al. clock^4^ is calculated from weighted methylation values of 8 CpGs, that were selected as the most informative CpGs in a training set of whole blood samples from 390 healthy individuals from the United Kingdom Ovarian Cancer Study, Human Aging-associated DNA Hypermethylation at Bivalent Chromatin Domains dataset, and Genome-wide Analysis of Autosomal Sex Differences in Human DNA Methylome dataset. The training population for this signature included individuals aged 20-78 with a mean age of 61.2. Importantly, 96.7% of the training set participants were female. This clock targeted older adults to calibrate DNAm age more accurately among adults compared to pre-adolescents. Results were not significantly influenced by sex, smoking, or variation in blood cell subpopulations. The Vidal-Bralo signature correlates with Horvath 1, Hannum, and Weidner signatures.

#### Zhang2019

The Zhang epigenetic clock was constructed from 12,661 blood and saliva samples from the Lothian Birth Cohort of 1921 and 1936 Study.^5^ The training set population for this signature had a mean age of 86 (Wave 3) when blood was collected. The signature is comprised of 514 CpG probes identified by an elastic net regression on chronological age. While this clock predicts a biological age that correlates with chronological age, it does not correlate with all-cause mortality in validations using subsets of the training population.

### Second Generation Clocks

#### Lin

The Lin epigenetic clock measures a relative risk of all-cause mortality.^6^ This 99 CpG model was originally trained on DNAm profiles of normal blood samples (n=446) with mortality data from the Lothian Birth Cohort 1921 Study. The mean age for participants in the training dataset was 79 years old and the authors note that it systematically overinflates the ages of younger samples. This signature correlates with chronological age and is associated with malignancy of tumor cells. This signature also correlates with telomere length.

#### PhenoAge

The PhenoAge clock predicts a phenotypic age trained on 9 clinical biomarkers (albumin, creatinine, serum glucose, CRP, lymphocyte percent, mean red cell volume, red cell distribution width, alkaline phosphatase, white blood cell count) which estimates an individual’s mortality risk.^7^ The PhenoAge clock is calculated from weighted methylation values at 513 CpGs in whole blood and was trained in the NHANES III study of individuals >=20 years old (n=9926). Levine et al. found that this clock correlated with age and predicted mortality better than first generation clocks.

PhenoAge predicts risk of multiple aging outcomes such as mortality, cancer, healthspan, physical function and Alzheimer’s disease and shows high correlation with biomarkers such as high CRP, glucose, triglycerides waist-to-hip ratio and low HDL cholesterol. PhenoAge was validated on data from several studies including NHANES IV, InCHIANTI, Jackson Heart Study, Women’s Health Initiative, Framingham Heart Study, and the Normative Aging Study.

#### GrimAge

GrimAge predicts a phenotypic age trained on 7 surrogates of plasma proteins and smoking pack years.^8^ First, authors defined surrogate biomarkers of physiological risk and stress factors with plasma proteins (including adrenomedullin, CRP, plasminogen activation inhibitor 1 (PAI-1) and growth differentiation factor 15 (GDF15)) and DNAm-based estimator of smoking pack-years. Then, time-to-death was regressed on these biomarkers and an estimator of smoking years to estimate a composite biomarker of lifespan. Lu et al. report that the rate of GrimAge-based aging has predictive ability for time to death, coronary heart disease, cancer and age-related conditions. The training data for this signature comes from the Framingham Heart Study including n=2356 individuals with mean ages of 66 years. 1030 CpGs associate with the 7 composite scores. This signature was validated on n=7375 participants from Inchiani, JHS, WHI and FHS. The signature is robust to adjustment for bmi, education, alcohol, smoking, diabetes, cancer, and hypertension. However, it fails to correlate with telomere length.

#### PC Clocks

There are many potential sources of technical variation in CpG beta values on which epigenetic clocks are based, leading to low reliability of CpG beta values (or M values), which can result in low intraclass correlation coefficients (ICCs) for epigenetic clock values in replicate samples. In an effort to increase the epigenetic aging signal to technical noise ratio of frequently used clocks, Higgins-Chen and colleagues retrained the Hannum, Horvath1, Horvath2, GrimAge, and PhenoAge clocks on PCs of CpG methylation values rather than on individual CpGs.^9^ This hypothetically increases the number of CpGs contributing to an epigenetic age calculation, thus reducing the effects of technical noise at any given individual CpG.

A unique set of methylation PCs was calculated for each clock, starting with a standard set of 78,464 CpGs that overlap across all original training datasets for the clocks and that are present on both the 450K and EPIC arrays (see supplementary table 6 of Higgins-Chen et al for details). PCA on the separate training datasets resulted in 655 PCs for Hannum, 4,280 PCs for Horvath1, 894 PCs for Horvath2, 3,934 PCs for GrimAge, and 4,504 PCs for PhenoAge. Elastic net regression was then used to train these PCs to predict either the original training target (for PCPhenoAge, which was trained on the same phenotypic age biomarker score used by Levine et al) or the original epigenetic clock age (for PC Hannum, PC Horvath 1, PC Horvath 2, and PC GrimAge) if complete training data from the original clock was not available. Thus, the PC versions of these clocks predict the same outcomes (chronological age, biological age, mortality, etc.) as their non-PC clocks. Elastic net regression retained 390 PCs for PCHannum, 121 PCs for PCHorvath1, 140 PCs for PCHorvath2, 1,936 PCs for PCGrimAge, and 652 PCs for PCPhenoAge. All PC clocks were subsequently validated in independent testing datasets and showed high correlation with the original clocks in training and testing datasets. The PC clocks were also tested on technical replicates and show improved ICC values compared to their original (non-PC) clocks for both clock ages and age acceleration (clock age regressed on chronological age).

PC Hannum, PCHorvath1, PCHorvath2, PCGrimAge, and PCPhenoAge were calculated in R using code available on GitHub at https://github.com/ MorganLevineLab/PC-Clocks/.

### Third Generation Clocks

#### DunedinPoAm

The DunedinPoAm epigenetic signature measures the pace of aging.^10^ This signature modeled the change over time of 18 biomarkers of organ system dysfunction in n=954 participants of the Dunedin Study. The age of the mostly White participants in this training set was 26 years old at the time of the first blood collection and 38 years at the time of the second blood collection. An elastic net regression was used to compute weights for 46 CpGs. The signature correlates modestly with Horvath, PhenoAge, and Hannum signatures but outperforms all of them as a proxy of self-rated health. The signature is Z-scaled such that the mean value of the analytical sample set is 0 and negative values indicate a reduced rate of aging compared to positive values.

#### DunedinPACE

The DunedinPACE epigenetic signature updates the DunedinPoAm epigenetic signature using the same approach, but this iteration includes 19 indicators of organ system integrity at 4 time points including a timepoint in which the oldest participants are 45 years old.^11^ DunedinPACE correlates with the DunedinPoAm (r = 0.57). This signature shows robust ability for replication. Like the DunedinPoAm signature, this signature is Z scaled such that negative values indicate slowed aging.

#### Zhang2017

Zhang et al. epigenetic signature is based on 10 CpGs that showed a strong association with all-cause mortality,^12^ which was selected from replicated results (58 out of 11,063 CpGs with FDR<0.05) from an epigenome-wide association study (EWAS) for all-cause mortality. This epigenetic signature specifically identifies those with increased risk of death by cancer and cardiovascular disease. The training set for this signature comes from the ESTHER study and includes 406 deceased participants with blood sampled at ages 50-75 years old. Although Zhang2017 may not strictly constitute a third generation of epigenetic clock, we group presentation of results for this clock with DunedinPoAm and DunedinPACE because, similar to those clocks, its unit of measurement is not epigenetic age in years, but rather, in this case, the risk of mortality.

### Sociodemographic and Lifestyle measures

Age at blood draw was calculated in years based on birth year (collected and validated across all waves of data) and calendar month and year of the in-home exam in Wave V when venous blood was drawn for methylation data. Sex assigned at birth is also cross-checked and validated across all Add Health waves of data. At Wave V Add Health used a combined race and ethnicity question in which participants self-identified their race and ethnicity and were permitted to check multiple categories. Those who selected multiple categories were then asked to select the race or ethnicity with which the participant most strongly identified. For a small proportion of cases (n=9) where Wave V information was missing or unavailable, Wave I was used to identify race and ethnicity. We included the following categories: White, Black, Hispanic, Asian/Pacific Islander, and “other” race (a combination of Native American and other). Immigrant generation was determined at Wave I based on the participant and their parent surveys. Generation 1 are those who were foreign-born with foreign-born parents; generation 2 are those who were U.S.-born with one or two foreign-born parents; and generation 3+ are those who were U.S.-born with both U.S.-born parents.

We used categorical responses to the Wave V survey question on the highest level of education and collapsed further into three categories of college or more; some college; and no college. Participants were asked what their total household income was before taxes and deductions in the last calendar year for all household members who contribute to household expenses; responses categories were provided in 13 brackets to reduce non-response. We further collapsed categories into the following four income levels: over $100,000; $49,999-$100,000; $24,999-$50,000; and $25,000 or less. The census region in which the participant lived at Wave V was coded from their address (Northeast, West, Midwest, South). Rural/urban residence patterns were derived from Wave V Contextual data based on the “Rural Urban Commuting Area” (RUCA) codes from 2010. Three mutually exclusive categories were constructed from participants’ description of the area in which their residence was located: metropolitan; micropolitan, small town or rural.

We constructed our measure of obesity status based on body mass index (BMI) constructed from measure height and weight at Wave V in the in-home exam. At Wave V, field staff measured height in cm from shoeless participants standing on uncarpeted floors and recorded weight to the nearest 0.1 kg. BMI was computed as kg/m^2^ and categorized obesity status as normal or underweight (BMI<25); overweight (25≤BMI<30), obese (30≤BMI<40), and severely obese (BMI≥40).^13,14^

Bouts of exercise per week were determined from five items in the Wave V survey that inquired about the number of times in the past week the participant performed the following forms of exercise respectively, aerobic activities, bicycling, gym activities, individual sports, or golf. The number of each of these types of activity was summed and categorized as: 0, 1-4 times per week, or 5+ times per week. At Wave V participants were asked whether they had ever smoked and whether they were current smokers. From these questions we categorized tobacco use as never; former and current. To categorize participants according to their usage of alcohol at Wave V, we first used the question whether they had ever drank alcohol, and if they answered that they had not, they were categorized as “None”. If participants said that they had ever drank, we then used questions on the number of days drank last month, days drank last year, and frequency of binge-drinking. If the participant engaged in binge drinking in the last year or had reported drinking daily in the last month or last year, they were categorized as “Heavy/Binge” and all other participants that drank less than daily and did not binge drink were categorized as “Light”.

---

**Supplement Table 1.**
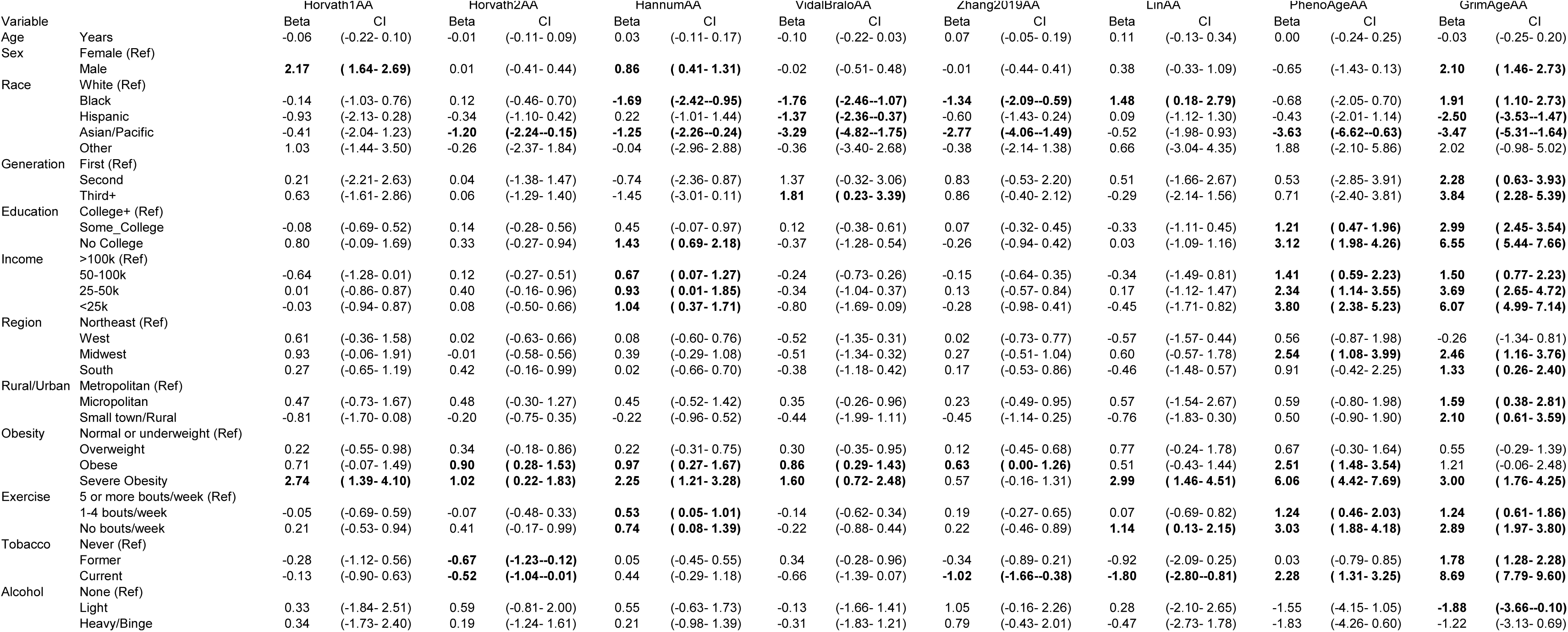
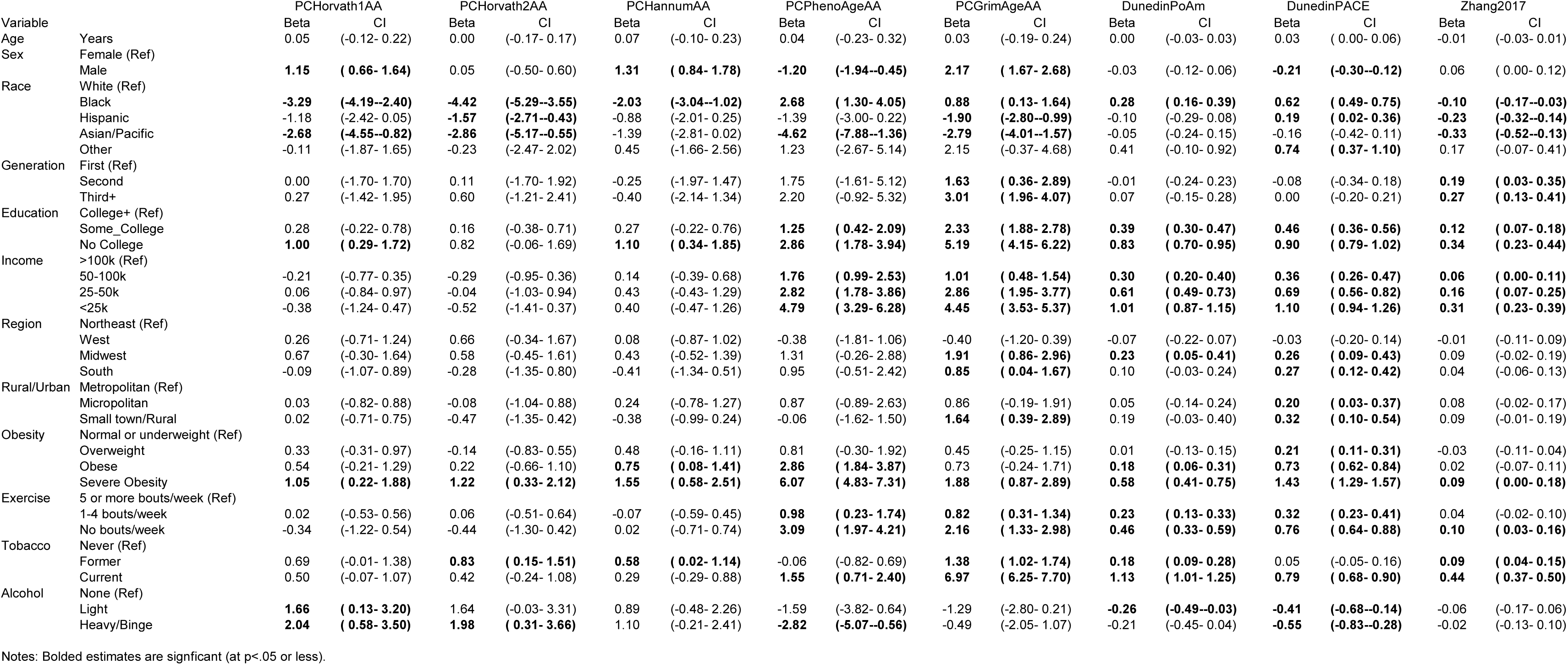
Epigenetic clock estimates from weighted bivariate models of sociodemographic and lifestyle characteristics (N=4237).

**Supplement Table 2.**
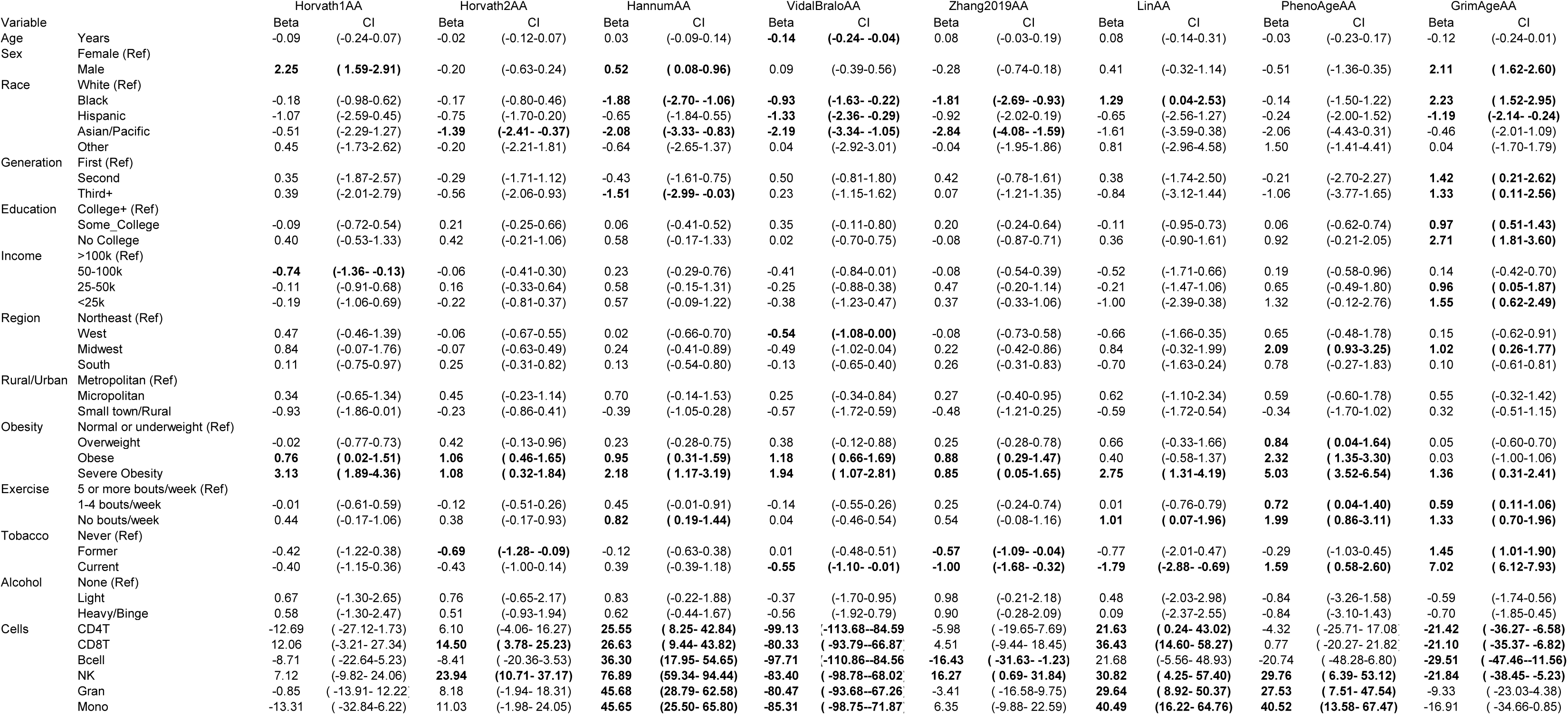
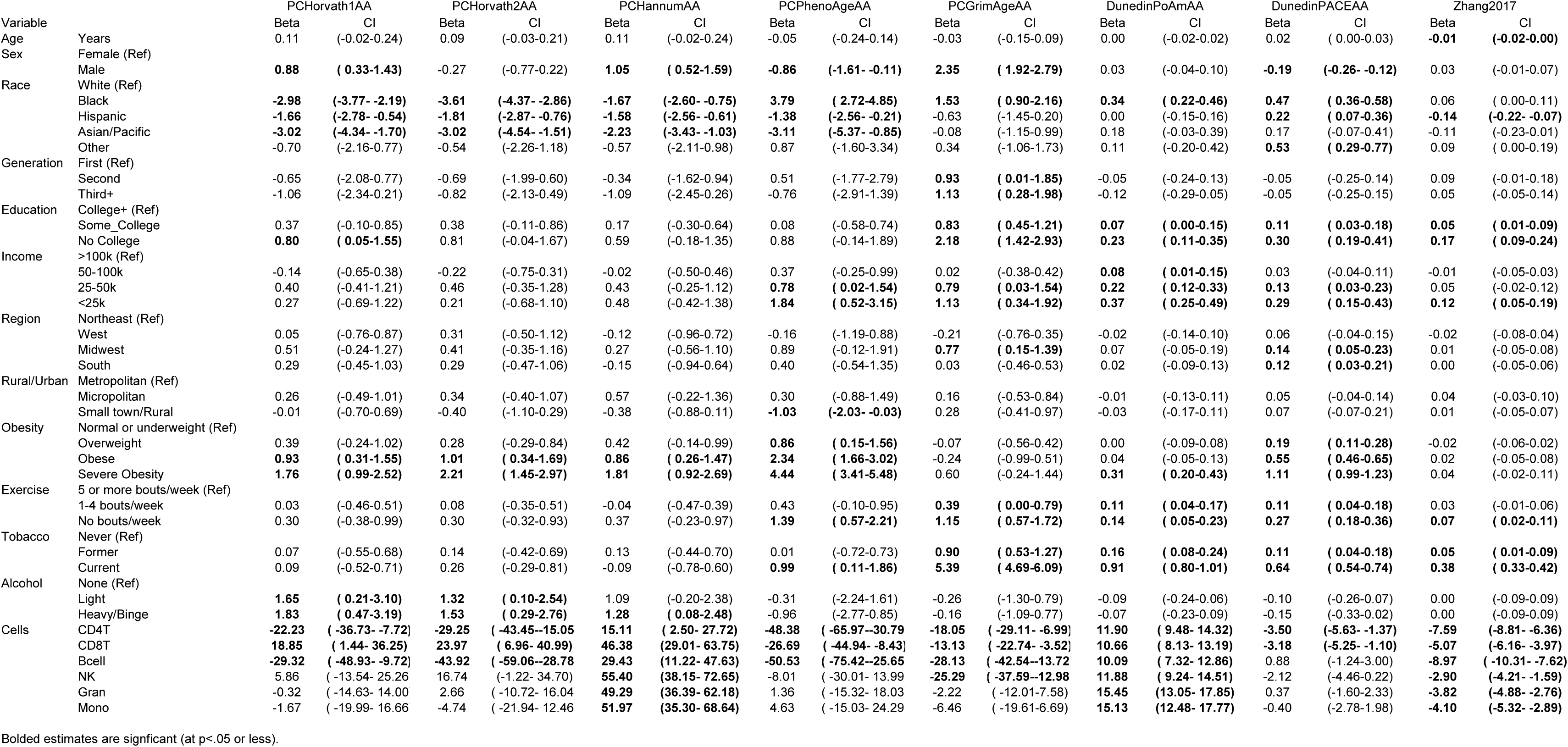
Epigenetic clock estimates from weighted multivariate models of sociodemographic and lifestyle characteristics, including cell composition (N=4237).

## Notes

### Competing Interest Statement

The authors have declared no competing interest.

### Summary of Updates

Orcid identifer for one author was corrected

